# *Anopheles (Kerteszia) cruzii*, the main malaria vector in the Brazilian Atlantic Forest, is a complex of at least five cryptic species

**DOI:** 10.1101/2025.11.14.688459

**Authors:** Kamila Voges, Guilherme de Rezende Dias, Eduardo Guimarães Dupim, André Nóbrega Pitaluga, Thyago Vanderlinde, Carlos José de Carvalho Pinto, Helder Ricas Rezende, Fabiana Uno, Sarah Jayne Forrester, James Chong, A. Bernardo Carvalho, Luísa D. P. Rona

**Author notes:** Equal senior authors (A. Bernardo Carvalho, Luísa D. P. Rona). E-mail addresses: GRD, EGD, ANP, TV, CJCP, HRR, FU, SJF, JC, ABC.

## Abstract

Malaria, a tropical disease caused by *Plasmodium* and transmitted by *Anopheles*, remains a public health concern in Brazil. While most cases occur in the Amazon, transmission persists in the Atlantic Forest, where *Anopheles* mosquitoes of the *Kerteszia* subgenus are the primary vectors of human and simian malaria. Previous studies using cytogenetics, isoenzymes, and molecular markers have suggested cryptic species within *Anopheles* (*Kerteszia*) *cruzii* and *Anopheles* (*Kerteszia*) *bellator*. We sequenced 55 genomes: 35 *An. cruzii* s.l. (four with Nanopore and 31 with Illumina), 12 *An. bellator* s.l., and eight *An. homunculus*, the latter two with Illumina. Phylogenomic analysis revealed at least five cryptic species within *An. cruzii* s.l., labelled A-E, with evidence of sympatry in some locations. *Anopheles bellator* s.l. also forms a species complex, comprising at least three distinct lineages. These cryptic species showed high genetic differentiation (*F_ST_* range: 0.4-0.7), typical of interspecific comparisons. In contrast, *An. homunculus* populations showed low differentiation (*F_ST_* ∼ 0.2), suggesting a single widespread species. Our analysis confirms cryptic speciation in *An. cruzii* and *An. bellator*, but not in *An. homunculus*. These findings are important for understanding malaria transmission in the Atlantic Forest, given that vector competence may differ among cryptic species.

## BACKGROUND

Malaria remains a major global public health challenge, affecting millions of people each year. According to the World Health Organization, there were 263 million cases worldwide in 2023^1^. In Brazil, most cases occur in the Amazon region; however, autochthonous cases have also been reported in the Atlantic Forest, particularly in the states of Rio de Janeiro, São Paulo, and Santa Catarina^2,3^. *Anopheles* (*Kerteszia*) *cruzii*, *Anopheles* (*Kerteszia*) *bellator*, and *Anopheles* (*Kerteszia*) *homunculus* are recognized as malaria vectors associated with bromeliads in the southern and southeastern regions of Brazil^4^. The most important vector in these regions is *An*. *cruzii*, a bromeliad-breeding mosquito known for transmitting both human and simian malaria^2,4–7^. This species ranges from Rio Grande do Sul in southern Brazil to Sergipe in the northeast and thrives in bromeliad-rich areas^8^, earning it the name “bromeliad malaria” vector^9,10^. In areas with large remnants of Atlantic Forest and abundant bromeliads (*e.g.,* Rio de Janeiro, São Paulo, Paraná, Santa Catarina), *An*. *cruzii* occurs at high densities, frequently enters houses, and feeds across vertical forest strata, from canopy to ground, biting both monkeys and humans^6,11,12^. This opportunistic behaviour enables natural infections with human (*Plasmodium vivax*, *Plasmodium falciparum*, *Plasmodium malariae*) and simian (*Plasmodium simium*, *Plasmodium brasilianum*) parasites^13–15^, with the practically indistinguishable *P. vivax / P. simium* responsible for most cases in the Atlantic Forest^3,16,17^. Consequently, autochthonous malaria in this region is a zoonosis^18^, and cross-transmission between primates and humans may occur^19^ in localities where infected monkeys (*e.g.,* howler monkeys, genus *Alouatta*) coexist with *An. cruzii*.

Numerous genetic studies have suggested that *An. cruzii* is a complex of cryptic species. For example, differences in *X* and 3*L* chromosomal inversion patterns among *An. cruzii* populations in southern and southeastern Brazil suggest the presence of distinct evolutionary units within this species^20,21^. In addition, isoenzyme analyses of *An. cruzii* populations from Santa Catarina, São Paulo, Rio de Janeiro, and Bahia revealed significant genetic divergence between the Bahia population (northeastern Brazil) and those from the South and Southeast^22^.

Studies using molecular markers have further confirmed the existence of multiple evolutionary lineages within the *An. cruzii* complex^23–27^. According to these studies, populations along the coastline and lower slopes of the Serra do Mar mountains (below 600 m) in southern and southeastern Brazil, spanning ∼1,000 km, likely belong to the same species, forming a major group within the *An. cruzii* complex. In contrast, the Bahia State population is genetically distinct, suggesting that it is a separate species within the complex^24,25^. Additionally, at least two cryptic species were identified at higher altitudes (above 900 m) in the Serra do Mar and Serra da Mantiqueira Mountain ranges. Notably, the Bocaina population (on the continental side of The Serra do Mar) contains two reproductively isolated groups living in sympatry^23–27^. In the *An. gambiae* s.l. species complex, sibling species differ in their ability to transmit malaria^28,29^. Similarly, the occurrence of more than one lineage of *An. cruzii* in the Atlantic Forest could have significant implications for malaria transmission and control in Brazil. Indeed, all historical and current autochthonous cases of human malaria in southern and southeastern Brazil (*e.g.,* Florianópolis and Guapimirim)^2,30,31^ align with the distribution of only one of the *An. cruzii* cryptic species identified through molecular studies (*e.g.,* using the *timeless* and *cpr* genes as markers or more than 2,000 genes, as in the present study)^23,27^. Therefore, exploring the population genomic structure of this species complex may help to clarify which lineages are relevant vectors of malaria.

Similarly, *An. bellator* appears to represent a cryptic species complex. Carvalho-Pinto and Lourenço-de-Oliveira^32^ used isoenzyme analysis to study populations of *An. bellator* from Florianópolis (Santa Catarina, southern Brazil), Cananéia (São Paulo, southeastern Brazil), and Itaparica (Bahia, northeastern Brazil). They found that populations from the South and Southeast are genetically more similar to each other than to the population from the Northeast. More recently, Voges et al.^33^ analyzed two molecular markers and also found strong genetic structuring among Brazilian populations of this species. Their results revealed the existence of two distinct groups within the Atlantic Forest: one in the South and Southeast regions of Brazil, and another in the Northeast (Camacan, Bahia). In contrast to the two previously mentioned species, available evidence suggests that *An. homunculus* does not represent a complex of cryptic species: Cardoso et al.^34^ analysed populations of *An. homunculus* from different regions of the Brazilian Atlantic Forest. Their results indicated that populations from Bahia (Northeast), Espírito Santo and São Paulo (Southeast), and Rio Grande do Sul (South) all seem to belong to the same species^34^.

Previous molecular studies of *An. cruzii*, *An. bellator*, and *An. homunculus* were based on a very small number of loci, which limits the scope and strength of their conclusions. Therefore, in the present study, we applied a phylogenomic approach to analyse genetic differentiation and infer the phylogenetic relationships among samples of these species collected across the Brazilian Atlantic Forest, aiming to provide a comprehensive assessment of their evolutionary and population structure.

## METHODOLOGY

### Mosquito collection

The mosquitoes used in this study were collected from various locations across the Brazilian Atlantic Forest (Figure 1): Itaparica (Bahia State-BA), Camacan (Bahia State-BA), Santa Teresa (Espírito Santo State-ES), Guapimirim (Rio de Janeiro State-RJ), Ilha Grande (RJ), Itatiaia (RJ), Bocaina (São Paulo State-SP), Campos do Jordão (SP), Paranapiacaba (SP), Santos (SP) and Florianópolis (Santa Catarina State-SC). The collection, treatment, and preservation of both adult and immature mosquitoes followed the methodology described by Dias et al.^27^. Detailed information about the samples, including geographical coordinates, sex, and life stage at the time of collection (adult or larvae), is provided in Additional File 1. Species identification was carried out following Consoli and Lourenço-de-Oliveira^8^ and Forattini^35^. Key morphological structures important for species-level identification were photographed for each individual using an Olympus SZX16 stereomicroscope. Following morphological identification, the mosquitoes were preserved in 100% ethanol at -20°C until DNA extraction.

**Figure 1.**
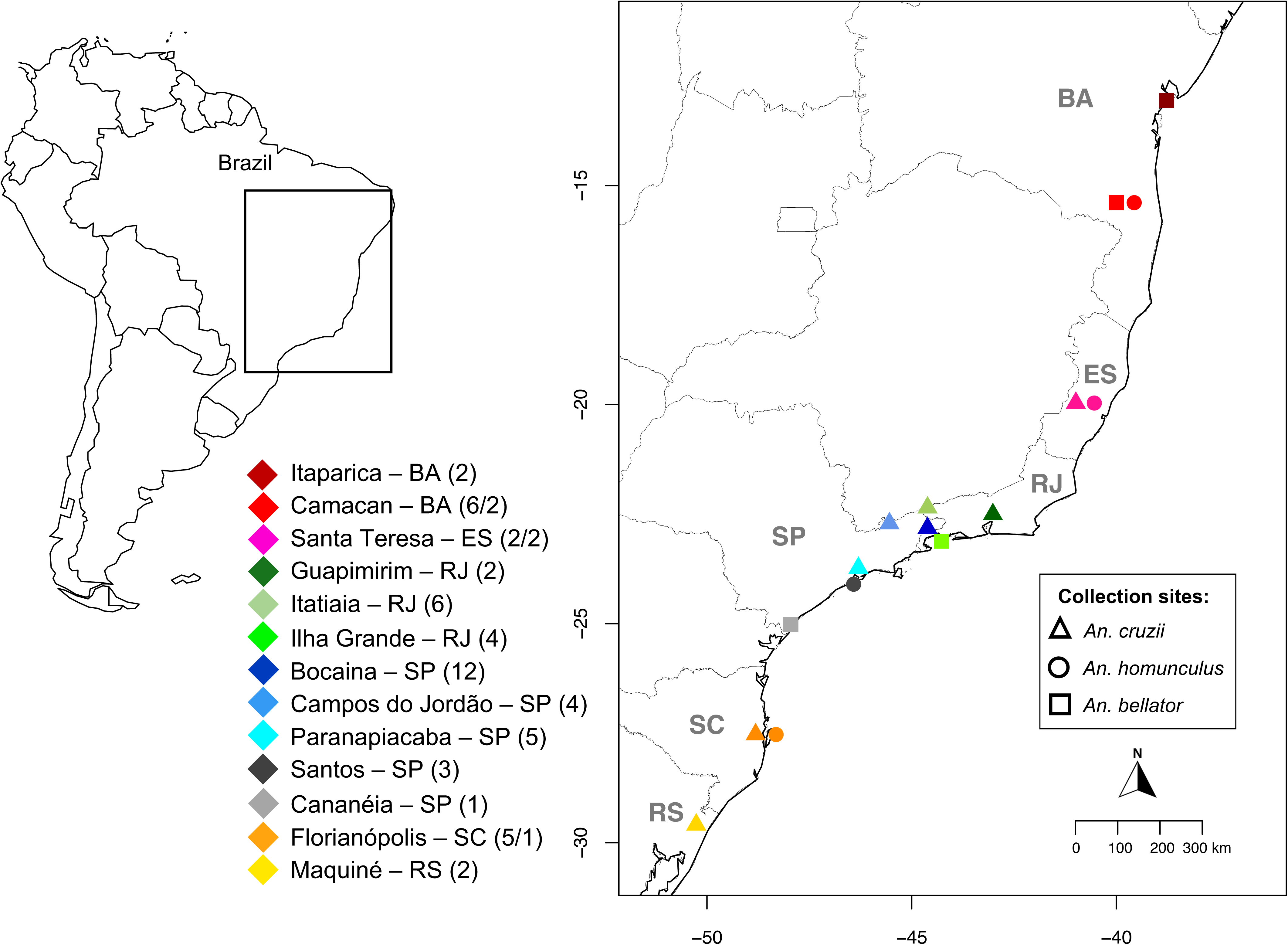
*Kerteszia* collection sites in the Brazilian Atlantic Forest. Left panel, map of South America. Right panel, a zoomed-in section showing the sample collection sites, marked with different symbols: triangles for *An. cruzii* s.l., circles for *An. homunculus*, and squares for *An. bellator*. The x- and y-axes of the zoomed-in map represent longitude and latitude, respectively. The number of samples collected from each location is indicated in parentheses. In Camacan, Santa Teresa, and Florianópolis, two *Kerteszia* species were captured; in these cases, the first symbol corresponds to the first number listed in brackets.Two genome samples of *An. cruzii s.l.* from Maquiné (GCA_943734635.1, GCA_964417045.1) and one *An. bellator* from Cananéia (GCA_943735745.1) were obtained from the Sanger *Anopheles* Reference Genomes Project.The maps were created using the *sf*, *maps*, *mapdata* and *rworldmap*packages^116–119^ in R Software, version 4.3.1^120^.

### DNA extraction

A non-destructive enzymatic method was used to extract DNA from individual mosquitoes, adapted from Santos et al.^36^. This approach preserves the morphology of taxonomically important structures, such as the exoskeleton and male genitalia. DNA extraction was carried out using the Qiagen Puregene Core Kit (cat # 158667). Mosquitoes were first split at the abdomen-thorax junction (to facilitate proteinase K digestion) and placed in individual tubes containing 100 µL of lysis solution and 1 µL of proteinase K (20 mg/mL; Qiagen, cat # 158918). Samples were incubated in this solution for three days at 45°C, followed by 1 minute on ice. Next, 33µl of precipitation solution was added to each sample, mixed by inversion, and incubated on ice for 5 minutes. The samples were then centrifuged for 3 minutes at 21,130 rcf, and the supernatant was transferred to a new tube while the old tube containing the pellet was discarded. Then, 0.5µl of RNase was added (4 ug/mL, Qiagen, cat # 158922), followed by three incubation steps: 15 minutes at 65°C, 15 minutes at 37°C, and a further 15 minutes at 65°C after adding 2µl of proteinase K. Samples were again placed on ice for 1 minute, 33µl of precipitation solution was added, mixed by inversion, and returned to ice for 5 minutes. After a second centrifugation at 21,130 rcf for 3 minutes, the supernatant was transferred to a new tube containing 1µl of Invitrogen™ GlycoBlue™ Coprecipitant (15 mg/mL, cat # AM9515), mixed, and then 100µl of absolute isopropanol was added and gently mixed by inversion. Samples were centrifuged at 21,130 rcf for 5 minutes, the supernatant was discarded, and the resulting blue pellets were briefly air-dried at room temperature. Each pellet was washed with 100µl of 70% ethanol, centrifuged at 21,130 rcf for 1 minute, and the supernatant discarded. Pellets were air-dried at room temperature for approximately 10 minutes. To dissolve the DNA, 50µl of DNA hydration solution was added to each tube, followed by incubation at 65°C for 1 hour; the tube was then left overnight at room temperature. DNA concentration was estimated using Qubit with dsDNA Quantitation High Sensitivity reagents (Invitrogen, cat # Q32851). These DNA samples were stored at -20°C until sequencing. The same procedures were applied to the *An. cruzii* Flo F3 N and *An. cruzii* Boc F3 N samples, before sequencing with Nanopore technology.

For the samples *An. cruzii* Boc F5 N and *An. cruzii* Boc M7 N, which were also sequenced using Nanopore technology, DNA was extracted from each mosquito using the protocol developed by Kim et al.^37^ to sequence single flies.

### Morphological characters of the male genitalia

The analysis of male genitalia, considered the most reliable method for morphological species identification in *Anopheles*^38^, was applied in this study. The protocol from Consoli & Lourenço-de-Oliveira^8^ was modified as follows: the male genitalia were carefully removed from the abdomen after DNA extraction and cleared in 10% KOH for 12 hours. The samples were then subjected to a dehydration process using a graded ethanol series: 70%, 80%, and 90% ethanol for 15 minutes each, followed by 95% and absolute ethanol for 10 minutes. To enhance visibility, a second clarification step was performed using xylene for 60 minutes. Finally, slides were mounted using Entellan (Sigma-Aldrich, cat # 107960) for analysis.

### Separation of the cryptic species in the Bocaina population

Dias et al.^27^ identified a potentially fixed difference in the *cpr* gene between two cryptic species of *An. cruzii* in the Bocaina population (referred to as “Bocaina Group 1” and “ Bocaina Group 2” by Dias et al.^27^; in the present study, we refer to them as species B and C, respectively). We exploited this difference to design a PCR assay for species identification. Namely, two species-specific forward primers were designed: Boc1F (5’GTGTAATATGGTAAGCGAACG) and Boc2F (5’GTGTAATATGGTAAGCGAAACG). These primers should work respectively only in *An. cruzii* species B and *An. cruzii* species C, when combined with a shared reverse primer BocR (5’TTTCTCGATGTCTTTCAGCT).

PCR amplifications were performed using an Applied Biosystems Veriti 96-Well Thermal Cycler (Model 9902) with GoTaq® Hot Start Polymerase (Promega, cat # M500B) under the following conditions: an initial denaturation step at 95°C for 9 minutes, followed by 40 cycles of 90°C for 30 seconds, 62°C for 30 seconds, and 72°C for 20 seconds, with a final extension at 72°C for 7 minutes. PCR products were applied to agarose gels (1%), stained with Ethidium Bromide, and photographed in a UV transilluminator. It is important to note that for sequenced samples (*e.g., An. cruzii* Boc F5 N, *An. cruzii* Boc M7 N, and *An. cruzii* Boc F4) this PCR assay was not necessary, as the species could be directly identified via phylogenomic analysis.

### Sequencing, genome assembly, and removal of contaminants

We sequenced a total of 55 samples: 51 using Illumina (SRA accessions on NCBI: SAMN48789258–SAMN48789358, BioProject: PRJNA1269491) and four using Nanopore (SRA accessions on NCBI: SAMN48793262–SAMN48793265, BioProject: PRJNA1269491). Illumina 150 bp paired-end libraries were prepared using the TruSeq Nano DNA or Nextera XT protocols (insert size: 350bp) and sequenced on a HiSeq 2000 at Macrogen, Korea (Additional File 1). To select the most suitable Illumina assembler for our dataset, seven *Anopheles* samples were assembled using both SPAdes-3.12.0^39^ (-t 100, -m 800, and default values for the other parameters) and Platanus 1.2.4^40^ (-t 32, -m 128, and default values for the other parameters). Assembly quality was evaluated using BUSCO v3^41^ (using the default parameters) based on the presence of orthologues from the Diptera reference set (odb9, downloaded from https://BUSCO.ezlab.org/, comprising 2,799 genes). The Platanus assembler, which demonstrated superior performance, was applied to the remaining Illumina samples.

We used Blobtools^42^ (with -l 200, --noreads, -r superkingdom, and default values for the other parameters) to identify and remove contaminant contigs from the primary assemblies, excluding all bacterial sequences from the final dataset. The tool requires a hits file and a coverage file: hits were generated via BLASTX and BLASTN searches against reference databases and filtered to remove human sequences and taxa on a contaminant ID list. The best hits per region were selected using the script *minusK_blob.awk* (k=3). Both SPAdes and Platanus produce FASTA files with deflines containing coverage information; we parsed them to generate a tab-delimited coverage file (scaffold name, length, coverage) compatible with Blobtools.

The final completeness of all samples was assessed using BUSCO v4.1.4^43^ with the Diptera reference set (odb10, downloaded from https://BUSCO.ezlab.org/, comprising 3,285 genes). Assembly quality metrics, such as N50, were estimated with QUAST^44^, using the default parameters. For Illumina samples, genome sizes were calculated from raw read coverage using Jellyfish-2.3.0^45^ and GenomeScope^46^, using -m 21 and the default parameters.

Three individuals (two from Florianópolis and one from Bocaina) were sequenced using Nanopore 9.4 flow cells on a MinION Mk1C. To assess the performance of different Nanopore assemblers, both Canu 2.1.1^47^ (genomeSize=200m, maxInputCoverage=100, and default values for the other parameters) and Flye 2.9.3-b1797^48^ (default parameters) were used. For the final assembly of these samples, we opted for Flye. The Florianópolis samples had low coverage (mean coverage of 6 and 17), resulting in low assembly quality (*e.g.*, N50 values of 9.269 kb and 294.561 kb). To address this, reads from these two individuals were combined and reassembled following the same procedures. This combined assembly was labelled as *An. cruzii* Flo F3 N, while the Bocaina sample was labelled as *An. cruzii* Boc F3 N. To enhance accuracy, three rounds of polishing were performed using Pilon v1.24^49^ with corresponding Illumina reads^40^. For *An. cruzii* Boc F3 N, Illumina reads from *An. cruzii* Boc F2, *An. cruzii* Boc M3, and *An. cruzii* Boc M2 were used; for *An. cruzii* Flo F3 N, reads from *An. cruzii* Flo M2, *An. cruzii* Flo M1, *An. cruzii* Flo M3, and *An. cruzii* Flo F2 were utilised. The polished assemblies were evaluated using BUSCO v4.1.4^43^ and QUAST^44^.

Two additional samples, *An. cruzii* Boc F5 N and *An. cruzii* Boc M7 N, were sequenced using Nanopore R10.4.1 flow cells on a PromethION2 system. Library preparation steps, including DNA repair, end preparation, bead clean-ups, and adapter ligation, followed the manufacturer’s protocols with adaptations from Kim et al.^37^. Base calling was performed using Dorado v0.7.1, applying a minimum quality score threshold of 10. Adapter sequences were removed using Porechop_ABI v0.5.0^50^, with the --ab_initio option. The cleaned reads were assembled using two different approaches to evaluate assembler performance: (i) with Flye v2.9.3-b1797^48^, using the raw reads without prior correction, and (ii) with Hifiasm v0.19.9-r616^51^, using reads previously corrected with HERRO^52^, as implemented in the ‘dorado correct’ function (Dorado v0.7.1, model v5.0.0 for R10.4.1/E8.2, default parameters). Assembly quality was again assessed using BUSCO v4.1.4^43^ and QUAST^44^. Finally, the assemblies were screened for contaminant sequences using NCBI Foreign Contamination Screen^53^.

In addition to the 55 genomes sequenced in this study, four reference genomes from the *Anopheles* Reference Genomes Project were included: GenBank assembly accessions GCA_943734635.1, GCA_964417045.1 (both *An. cruzii* specimens collected in Maquiné–RS), GCA_964417115.1 (*An. cruzii* collected in Itatiaia–RJ), and GCA_943735745.2 (*An. bellator* collected in Cananéia–SP).

### Search for orthologues and gene annotation

BUSCO v4.1.4 was used to identify single-copy orthologues in the 59 assemblies mentioned above, targeting 3,285 genes conserved among Diptera from the OrthoDB database (odb10, downloaded from https://BUSCO.ezlab.org/). These single-copy orthologues were employed for phylogenomic analysis, following the methodology described by Waterhouse et al.^54^, with modifications introduced by Dias et al.^55^. Briefly, the annotated orthologues from BUSCO v4.1.4 were processed using the scripts *fix_busco_CDS_frame.awk*^55^ and *BUSCO_cleaning_pipeline.awk*^55^ which correct annotation errors that could compromise the accuracy of phylogenomic analyses. Nucleotide sequences were aligned based on their encoded protein sequences using the *translatorex_vLocal.pl* script^56^.

### Phylogenomic inferences

Although BUSCO v4.1.4 aims to annotate all 3,285 genes from the odb10 database, some genes are inevitably absent from some assemblies. Previous studies have shown that including a large number of genes, even when not all are present in every sample, yields better performance in phylogenomic analyses than using a smaller subset of genes present across all taxa^57,58^. These findings are reassuring, suggesting that missing data does not pose a major issue in our phylogenomics analyses.

Nevertheless, in order to ensure that our conclusions were not compromised by artefacts arising from missing data, we conducted all analyses twice, employing both stringent and relaxed approaches. The stringent approach included only genes present in at least 50 of the 59 samples, thereby providing a more balanced dataset and reducing biases in species tree estimation^59^. However, this approach results in fewer genes, potentially reducing the statistical power. Conversely, the relaxed approach included genes present in at least 30 of the 59 samples, yielding a larger dataset but with a greater potential for bias. The stringent and the relaxed datasets had respectively 2,480 and 3,098 orthologues. We also considered using an even more stringent dataset by requiring the presence of a gene in all 59 samples. However, this resulted in too few genes (62).

Maximum-likelihood phylogenies were inferred for each gene in both datasets using the best-fit models identified by IQ-Tree3.0.0^60^. To address potential biases introduced by outlier sequences with long branches, TreeShrink^61^ was applied to the gene trees generated by IQ-Tree. These outliers, possibly resulting from annotation errors (*e.g.,* misidentification of paralogues as orthologues) or biological factors (*e.g.,* genes undergoing accelerated evolution), were removed to enhance analysis accuracy. Following outlier removal, species trees were inferred from both datasets using two approaches: maximum-likelihood (ML) and multispecies coalescent (MSC). The “filtered” alignments generated by TreeShrink, excluding problematic sequences, were concatenated and used for ML inference in IQ-Tree3.0.0^60^. The “pruned” gene trees, with outliers removed, were used for MSC inference in ASTRAL^62^, under default settings.

Sequences from *An. bellator* and *An. humunculus* were included to root the resulting trees, as these species are recognized as distinct from *An. cruzii* but belonging to the same subgenus. The command lines used for the phylogenomic inferences are available on GitHub (https://github.com/kamilavoges/Phylogenomics_Kerteszia/blob/main/phylogenomic_in ferences.txt).

### Genetic differentiation

Raw Illumina reads from each sample were normalised to 30× coverage using *seqtk*^63^ (version 1.3-r106) and then aligned to the reference genomes using *bwa*^64^ (version 0.7.17-r1194-dirty) and *samtools* 1.9^65^. The reference genomes came from the “*Anopheles* Reference Genomes Project” (https://www.sanger.ac.uk/collaboration/anopheles-reference-genomes-project/), which provided annotated genomes. Specifically, *An. cruzii* s.l. reads were aligned to the *An. cruzii* reference genome (accession GCA_943734635.1), while *An. bellator* and *An. homunculus* reads were aligned to the *An. bellator* reference genome (accession GCA_943735745.2). BAM files were subsequently processed with *Picard* v2.18.22^66^. SNP calling was performed on the processed BAM files using *freebayes* v1.0.2^67^, incorporating a CNV.bed file to distinguish male and female samples. Individual VCF files were generated for each sample and then merged into a single file using *BCFtools* v1.18^68^. The merged VCF was split by chromosome using *VCFtools* v0.1.17^69^, and missing genotypes were corrected using fixvcfmissinggenotypes.jar v3d336e5 ^70^.

*F_ST_* analyses were carried out using the VCF files and the *snpgds Fst* function from the R SNPRelate SNPRelate 1.42.0 package to quantify genetic differentiation between populations. *F_ST_* graphs were generated using R version 4.4.1 and custom scripts.

Genetic variation among *An*. *cruzii* s.l. samples was also assessed using Principal Component Analysis (PCA)^71^, implemented in scikit-allel^72^ via allel.pca (gn, n_components = 10, scaler = ‘patterson’, default value for the other parameters) on 20,319,109 biallelic SNPs from the whole genome. Individual ancestry, population structure, and admixture were inferred using the maximum likelihood method implemented in ADMIXTURE v1.3.0^73^. We used plink2 (v2.0.0-a.6.20LM) to filter the VCF file and convert it to the bed format required by ADMIXTURE, with the following parameters: --vcf-half-call m --max-alleles 2 --maf 0.01 --geno 0.10 --var-min-qual 30. Results were visualized using the custom python script plot_ADMIXTURE_v0.py.

The scripts used for all steps are available at https://github.com/kamilavoges/Phylogenomics_Kerteszia/blob/main/VCF_pipeline_an d_Fst_calculation.bash.

### Divergence times

We estimated divergence times with the RelTime method^74^ using a maximum likelihood tree inferred from *An. cruzii* s.l. sequences in IQ-Tree^60^. We used all codon positions to build the tree, but only third positions, assumed to evolve under near-neutral rates, were used for dating. *Anopheles bellator* and *An. homunculus* served as outgroups, and analyses were conducted in MEGA11^74^.

RelTime estimates divergence times from substitution rates. Since rates are unavailable for *An. cruzii* and RelTime does not account for calibration uncertainty, we tested four alternatives: (i) 19.0 substitutions/kbp/Myr, based on drosophilid colonization of the Hawaiian Islands^75^; (ii) 16.7 substitutions/kbp/Myr, estimated for drosophilid 3^rd^ codon positions from fossil calibration (Dias et al., *in press*); and (iii) 13.6 substitutions/kbp/Myr and (iv) 27.2 substitutions/kbp/Myr, derived from the spontaneous mutation rate of *Anopheles stephensi* (1.36 ×10^-9^ substitutions/site/generation)^76^, assuming 10 and 20 generations per year, respectively. This conversion was made to express the mutation rate in comparable units (substitutions/kbp/Myr). Drosophilid rates were included given their widespread use in Diptera molecular dating (*e.g.,* references 75,77)^75,77^, while *An. stephensi* provides the closest calibration within *Anopheles*^76^.

## RESULTS

### Assembly and quality of genomes

Our study presents the first draft genomes for several *Kerteszia* species, including *An. homunculus* and different species within the *An. cruzii* s.l. and *An. bellator* s.l. complexes. A total of 55 genomes were sequenced from nine populations: 35 samples from *An. cruzii* s.l., 12 from *An. bellator* complex, and 8 from *An. homunculus*. In addition, four genomes from the Sanger *Anopheles* Reference Genomes Project were included: three from *An. cruzii* s.l. and one from *An. bellator* s.l (Additional File 1). According to GenomeScope, Illumina genome coverages ranged from 32× to 141× (Additional File 1). GenomeScope also reported high levels of heterozygosity, averaging ∼ 2% (range: 1.3% - 3%), which is expected in field-collected insect samples (Additional File 1).

As a preliminary test, we assembled seven Illumina datasets using Platanus and SPAdes; Platanus outperformed SPAdes in all samples, with greater than 20% increase in complete orthologues (Additional File 2). This outcome is expected, given that Platanus was designed to handle highly heterozygous genomes (> 1%)^40^, while SPAdes was initially developed for smaller genomes, such as those of bacteria^39^. Consequently, Platanus was used for all 51 Illumina-sequenced genomes in this study. Based on GenomeScope analyses of the raw reads, the estimated average genome sizes were: 156 Mb (148 - 172 Mb) for *An. homunculus*, 170 Mb (154 - 181 Mb) for *An. cruzii* s.l., and 174 Mb (162 - 177 Mb) for *An. bellator*. These values are similar to that of *Anopheles* (*Nyssorhynchus*) *darlingi* (182 Mb)^78^ but smaller than that of *Anopheles* (*Cellia*) *gambiae* (278 Mb)^79^. They are larger than genome sizes calculated from assembly data, likely due to the collapse of repetitive regions during the assembly of Illumina short reads (Additional File 1). As expected, samples sequenced with Nanopore long-reads (and assembled with Flye or hifiasm) yielded genome sizes more consistent with GenomeScope estimates, *e.g.* 170 Mb for *An. cruzii* Flo F3 N and 183 Mb for *An. cruzii* Boc F3 N (Additional File 1).

Of the 46 samples sequenced using Illumina TruSeq, BUSCO v4.1.4 analysis showed that 28 had ≥ 90% complete orthologues (Additional File 1). N50 values ranged from 4 to 88 kb, with the largest contig size reaching 612 kb (Additional File 1). In contrast, the five samples sequenced with Illumina Nextera XT yielded assemblies of substantially lower quality: N50 values ranged from 3 to 8kb, with a maximum contig size of 143 kb, and BUSCO v4.1.4 values ranged from 36% to 67% of complete orthologue sequences. Nextera XT was used in cases where DNA quantities were below the recommended minimum for the Illumina TruSeq protocol (100 ng); this limitation, coupled with known biases in the Nextera XT protocol^80^ likely explains the reduced quality of these assemblies. Finally, as expected, the Nanopore assemblies were by far the best ones: N50 values ranged from 386 kb to 16.8 Mbp, with maximum contig sizes of up to 34 Mbp, and all BUSCO completeness scores exceeded 98.5% (see Additional File 1 and Additional File 3 for more details).

As frequently occurs in genome assemblies, we detected contaminant sequences using the Blobtools pipeline and removed them prior to analysis. The most common contaminants were bacteria (in decreasing abundance: *Proteobacteria*, *Actinobacteria*, *Bacteroidetes*, *Firmicutes*), with quantities varying widely across samples, from a few hundred base pairs to a few million, with a typical value of around 200 kb. Fungi (*Ascomycota*, *Basidiomycota*) were also fairly common. The most likely source of these contaminants is the insects’ gut.

### Differentiation of An. cruzii B and An. cruzii C in the Bocaina population

Dias et al.^27^ identified by sequencing two distinct *cpr* gene haplotypes in a sample of 12 *An. cruzii* s.l. from Bocaina – SP, along with a significant heterozygote deficit, suggesting cryptic speciation. We extended that study using a larger sample of 145 wild-caught mosquitoes from Bocaina and genotyping them with allele-specific PCR primers for the *cpr* gene (Figure 2).

**Figure 2.**
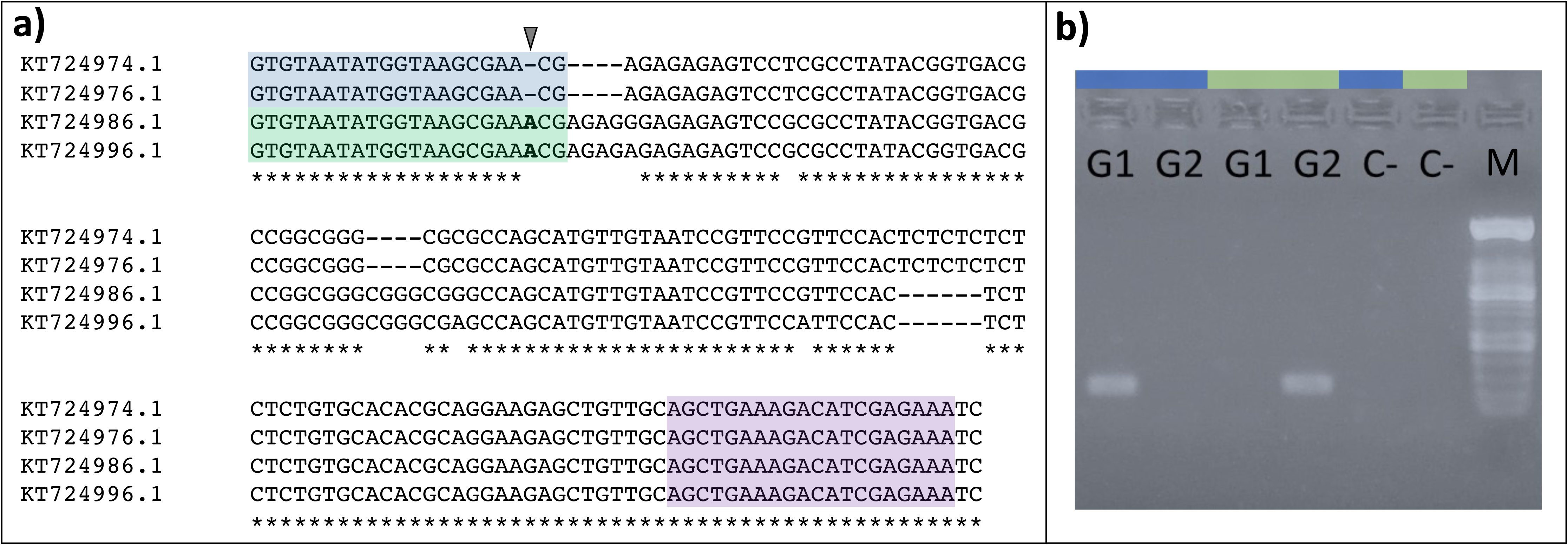
Identification of *Anopheles cruzii* B and C from Bocaina. **a)** Alignment of the *cpr* gene fragment in *An. cruzii* s.l.: Dias et al. (2018b) identified a likely fixed difference in the *cpr* gene between two cryptic species within the Bocaina population. They referred to these as “Bocaina Group 1” (accession numbers: KT724974.1 and KT724976.1) and “Bocaina Group 2” (accession numbers: KT724986.1 and KT724996.1). In the present study, we refer to these groups as *An. cruzii* B and C, respectively. In order to differentiate them, two species-specific forward primers were designed: Boc1F (5’-GTGTAATATGGTAAGCGAACG, in blue) for *An. cruzii* B, and Boc2F (5’-GTGTAATATGGTAAGCGAAACG, in green) for *An. cruzii* C. Each primer amplified only its respective species when paired with a common reverse primer, BocR (5’-TTTCTCGATGTCTTTCAGCT, in purple). The inverted triangle marks the position of the single nucleotide difference between the two forward primers. **b)** Agarose gel electrophoresis (1%) showing samples classified into *An. cruzii* B (G1) and *An. cruzii* C (G2). In the samples highlighted in blue, the primers Boc1F and BocR were used, while those highlighted in green used the Boc2F and BocR primers. The results confirm the specificity of the designed primers: G1 samples were amplified exclusively with the Boc1F primer, specific to *An. cruzii* B, whereas G2 samples were amplified only with the Boc2F primer, specific to *An. cruzii* C. Both primer sets produced an ∼180 bp fragment of the *cpr* gene. C-: negative controls (water instead of DNA). M: 100 bp molecular marker.

As the *cpr* gene is X-linked in *An. cruzii* s.l. (according to the reference assembly GCA_943734635.1), sex must be considered when calculating allele frequencies. Based on the PCR results, among the 145 genotyped mosquitoes, four males were *cpr^B^*/Y, three females were *cpr^B^*/*cpr^B^*, 54 males were *cpr^C^*/Y, and 84 females were *cpr^C^*/*cpr^C^*. No heterozygous *cpr^B^*/*cpr^C^* females were observed. The *cpr^B^* allele frequency in females was 3.45%, and under Hardy-Weinberg equilibrium, we would expect to find the following female counts: *cpr^B^*/*cpr^B^*, 0.103; *cpr^B^*/*cpr^C^*, 5.793; *cpr^C^*/*cpr^C^*, 81.103. The deviation from Hardy-Weinberg equilibrium is highly significant (*P* < 10^-5^, Fisher’s Exact Test^81^) with a complete absence of heterozygotes. The most plausible explanation for this heterozygote deficiency in sympatry is reproductive isolation, in this case, without any overt morphological diagnostic character (*i.e.,* cryptic speciation). The cryptic speciation hypothesis was further supported by whole-genome sequencing of four individuals carrying the *cpr^B^* allele (two males and two females) and seven carrying the *cpr^C^* allele (five males and two females), which revealed strong genetic differentiation across the genome, as detailed in the following sections.

Additionally, phylogenomic analyses (below) identified a third genetically distinct species in Bocaina, provisionally labelled *An. cruzii* D. This third cryptic species was represented by a single female (*An. cruzii* Boc F4), which was not genotyped using the *cpr* marker. However, its *cpr* sequence was retrieved from the assembled genome and showed several differences when compared with the sympatric *An. cruzii* B and *An. cruzii* C (Additional File 4).

### Phylogenomic inferences

Species trees were inferred for 59 samples of *An. cruzii* s.l. and related species (*An. bellator* and *An. homunculus*) using both maximum likelihood and multispecies coalescent methods. These analyses were performed with (*i*) 2,480 orthologues present in at least 50 of the 59 samples (the stringent dataset; data not shown), and (*ii*) 3,098 orthologues present in at least 30 of the 59 samples (the relaxed dataset). All four approaches yielded identical topologies; Figure 3 presents the results of the multispecies coalescent using the relaxed dataset.

**Figure 3.**
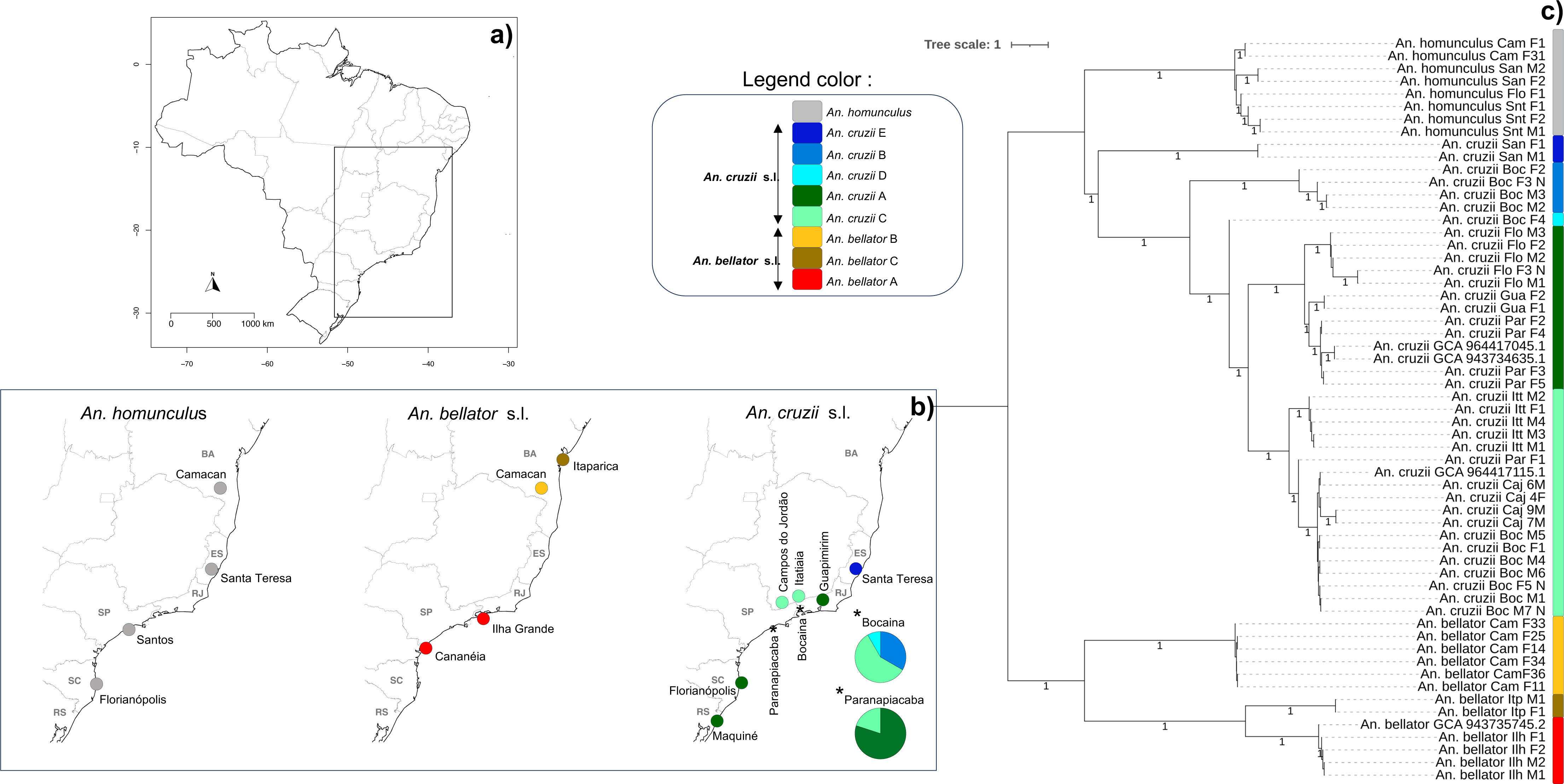
Cryptic speciation in *Anopheles cruzii* and *Anopheles bellator*. Map of Brazil **(a)** with a zoomed-in view of the boxed area **(b)**, showing the *Kerteszia* collection sites within the Atlantic Forest. **c)** Species tree inferred with Multispecies Coalescent (MSC) approach from 3,098 genes present in at least 30 of the 59 *Kerteszia* samples. The phylogenomic analysis shows no clear separation among *An. homunculus* populations (highlighted in grey). In contrast, cryptic species are present within *An. bellator*, which comprises at least three species (represented by shades ranging from reddish to yellowish), and within *An. cruzii*, which consists of at least five cryptic species (represented by the green and blue colors). Two genome samples of *An. cruzii s.l.* from Maquiné (GCA_943734635.1, GCA_964417045.1), one *An. cruzii s.l.* from Itatiaia (GCA_964417115.1), and one *An. bellator* from Cananéia (GCA_943735745.1) were obtained from the Sanger *Anopheles* Reference Genomes Project. Itp: Itaparica; Cam: Camacan; San: Santa Teresa; Gua: Guapimirim; Itt: Itatiaia; Ilh: Ilha Grande; Boc: Bocaina; Caj: Campos do Jordão; Par: Paranapiacaba; Snt: Santos; Flo: Florianópolis. M: males; F: females; N: Nanopore. The maps were created using the *sf*, *maps*, and *mapdata* packages^116–119^ in R Software, version 4.3.1^120^.

The phylogenomic analyses strongly and consistently support that *An. cruzii* s.l. comprises at least five cryptic species. One widely distributed group, provisionally labelled *An. cruzii* A, includes populations from the South and Southeast regions of Brazil, located in the Serra do Mar coastal zone. This group is represented by samples from Maquiné – RS, Florianópolis – SC, Paranapiacaba – SP, and Guapimirim – RJ. *Anopheles cruzii* B and D were found only in Bocaina – SP. *Anopheles cruzii* C was present in both Serra do Mar and Serra da Mantiqueira, two parallel mountain ranges in Southeast/South Brazil (Bocaina – SP, Paranapiacaba – SP, Campos do Jordão – SP, and Itatiaia – RJ). Finally, *An. cruzii* E was detected only in Santa Teresa – ES. As shown below, this is the only species with a morphological diagnostic character.

Across southern to northeastern Brazil, *An. cruzii* s.l. coexists with *An. bellator* and *An. homunculus*^34,82^. Phylogenomic analyses revealed that geographically distant populations of *An. homunculus* from Florianópolis – SC, Santos – SP, Santa Teresa – ES, and Camacan – BA exhibit low genetic differentiation (as indicated by the short branches in Figure 3), suggesting that they all belong to the same species.

For *An. bellator*, the phylogenomic analysis indicates that it comprises at least three cryptic species. *An. bellator* A appears to be widespread in southeastern Brazil, including Cananéia – SP and Ilha Grande – RJ. The second and third species occur in northeastern Brazil: *An. bellator* B was found in Camacan – BA, and *An. bellator* C in Itaparica – BA (Figure 3). Despite Itaparica being geographically close to Camacan (∼300 km), its *An. bellator* population is phylogenetically closer to those from Ilha Grande and Cananéia (> 2,000 km). This again demonstrates that genetic differentiation in these *Kerteszia* species is not primarily driven by geographic distance.

### Genetic differentiation (F_ST_)

Pairwise *F_ST_* values were estimated for all populations sampled in this study. All pairwise comparisons between different species within *An. cruzii* s.l. exhibited high *F_ST_* values (Table 1, Additional File 5).

**Table 1.** Mean *F_ST_* values for comparisons among *An.cruzii* s.l. populations. Species were ascertained based on phylogenomic analysis and sympatry (see text). The number of genes analysed was 6,663 (all), 461 (ChrX), 2,644 (Chr2), and 3,558 (Chr3). The numbers of individuals per population is indicated in parentheses; note that, except for X-linked genes in males, the number of sampled alleles is twice the number of individuals. Population abbreviations: Florianópolis (Flo); Paranapiacaba (Par); Campos de Jordão (Caj); Bocaina (Boc); Itatiaia (Itt); Guapimirim (Gua); Santa Teresa (San).

| Species | Populations | All genes | ChrX | Chr2 | Chr3 |
| --- | --- | --- | --- | --- | --- |
| *An. cruzii* B × *An. cruzii* E | Boc (4), San (2) | 0.728 | 0.877 | 0.722 | 0.713 |
| *An. cruzii* D × *An. cruzii* E | Boc (1), San (2) | 0.705 | 0.868 | 0.689 | 0.696 |
| *An. cruzii* C × *An. cruzii* E | Par (1), San (2) | 0.701 | 0.848 | 0.687 | 0.692 |
| *An. cruzii* C × *An. cruzii* E | Caj (4), San (2) | 0.690 | 0.886 | 0.665 | 0.682 |
| *An. cruzii* A × *An. cruzii* E | Flo (5), San (2) | 0.681 | 0.896 | 0.670 | 0.662 |
| *An. cruzii* C × *An. cruzii* E | Boc (7), San (2) | 0.676 | 0.875 | 0.654 | 0.666 |
| *An. cruzii* A × *An. cruzii* E | Gua (2), San (2) | 0.675 | 0.881 | 0.659 | 0.661 |
| *An. cruzii* C × *An. cruzii* E | Itt (5), San (2) | 0.661 | 0.858 | 0.644 | 0.648 |
| *An. cruzii* A × *An. cruzii* E | Par (4), San (2) | 0.660 | 0.875 | 0.640 | 0.648 |
| *An. cruzii* B × *An. cruzii* D | Boc (4), Boc (1) | 0.624 | 0.867 | 0.623 | 0.593 |
| *An. cruzii* B × *An. cruzii* C | Boc (4), Par (1) | 0.617 | 0.840 | 0.619 | 0.586 |
| *An. cruzii* B × *An. cruzii* A | Boc (4), Gua (2) | 0.578 | 0.865 | 0.573 | 0.544 |
| *An. cruzii* B × *An. cruzii* C | Boc (4), Caj (4) | 0.575 | 0.851 | 0.558 | 0.553 |
| *An. cruzii* B × *An. cruzii* A | Boc (4), Flo (5) | 0.572 | 0.870 | 0.569 | 0.536 |
| *An. cruzii* B × *An. cruzii* C | Boc (4), Boc (7) | 0.554 | 0.830 | 0.540 | 0.528 |
| *An. cruzii* B × *An. cruzii* A | Boc (4), Par (4) | 0.546 | 0.840 | 0.535 | 0.516 |
| *An. cruzii* B × *An. cruzii* C | Boc (4), Itt (5) | 0.535 | 0.807 | 0.527 | 0.506 |
| *An. cruzii* D × *An. cruzii* C | Boc (1), Par (1) | 0.507 | 0.814 | 0.491 | 0.479 |
| *An. cruzii* D × *An. cruzii* A | Boc (1), Flo (5) | 0.495 | 0.873 | 0.486 | 0.453 |
| *An. cruzii* D × *An. cruzii* C | Boc (1), Caj (4) | 0.494 | 0.842 | 0.461 | 0.474 |
| *An. cruzii* D × *An. cruzii* A | Boc (1), Gua (2) | 0.475 | 0.876 | 0.455 | 0.437 |
| *An. cruzii* D × *An. cruzii* C | Boc (1), Boc (7) | 0.468 | 0.810 | 0.441 | 0.444 |
| *An. cruzii* D × *An. cruzii* A | Boc (1), Par (4) | 0.456 | 0.828 | 0.432 | 0.425 |
| *An. cruzii* D × *An. cruzii* C | Boc (1), Itt (5) | 0.439 | 0.775 | 0.421 | 0.409 |
| *An. cruzii* C × *An. cruzii* A | Par (1), Flo (5) | 0.432 | 0.835 | 0.415 | 0.392 |
| *An. cruzii* C × *An. cruzii* A | Caj (4), Flo (5) | 0.432 | 0.836 | 0.399 | 0.405 |
| *An. cruzii* C × *An. cruzii* A | Caj (4), Gua (2) | 0.420 | 0.829 | 0.377 | 0.399 |
| *An. cruzii* A × *An. cruzii* C | Flo (5), Boc (7) | 0.415 | 0.812 | 0.386 | 0.386 |
| *An. cruzii* C × *An. cruzii* A | Par (1), Gua (2) | 0.407 | 0.823 | 0.380 | 0.374 |
| *An. cruzii* C × *An. cruzii* A | Caj (4), Par (4) | 0.402 | 0.803 | 0.359 | 0.382 |
| *An. cruzii* A × *An. cruzii* C | Gua (2), Boc (7) | 0.399 | 0.800 | 0.362 | 0.374 |
| *An. cruzii* A × *An. cruzii* C | Par (4), Boc (7) | 0.387 | 0.782 | 0.349 | 0.363 |
| *An. cruzii* C × *An. cruzii* A | Par (1), Par (4) | 0.387 | 0.782 | 0.355 | 0.360 |
| *An. cruzii* C × *An. cruzii* A | Itt (5), Gua (2) | 0.375 | 0.766 | 0.350 | 0.344 |
| *An. cruzii* C × *An. cruzii* A | Itt (5), Par (4) | 0.367 | 0.754 | 0.339 | 0.338 |
| *An. cruzii* C × *An. cruzii* C | Par (1), Caj (4) | 0.247 | 0.477 | 0.209 | 0.245 |
| *An. cruzii* C × *An. cruzii* C | Par (1), Itt (5) | 0.242 | 0.454 | 0.225 | 0.227 |
| *An. cruzii* C × *An. cruzii* C | Caj (4), Itt (5) | 0.235 | 0.513 | 0.201 | 0.224 |
| *An. cruzii* C × *An. cruzii* C | Itt (5), Boc (7) | 0.219 | 0.487 | 0.190 | 0.206 |
| *An. cruzii* A × *An. cruzii* A | Gua (2), Flo (5) | 0.212 | 0.625 | 0.193 | 0.172 |
| *An. cruzii* C × *An. cruzii* C | Par (1), Boc (7) | 0.206 | 0.408 | 0.179 | 0.201 |
| *An. cruzii* A × *An. cruzii* A | Par (4), Flo (5) | 0.180 | 0.548 | 0.156 | 0.151 |
| *An. cruzii* A × *An. cruzii* A | Gua (2), Par (4) | 0.140 | 0.485 | 0.119 | 0.112 |
| *An. cruzii* C × *An. cruzii* C | Caj (4), Boc (7) | 0.052 | 0.185 | 0.041 | 0.042 |

The case of the Bocaina samples of *An. cruzii* B and *An. cruzii* C is particularly noteworthy. In this comparison, high *F_ST_* values (> 0.5, Table 1, Figure 4) were observed across all chromosomes, confirming that the *cpr* gene differentiation previously reported by Dias et al.^27^ corresponds to two cryptic species. Alternative explanations, such as genotyping error or selection against heterozygotes, were ruled out, as these processes cannot account for genome-wide differentiation. As shown in Table 1, the third cryptic species in Bocaina (*An. cruzii* D) also shows high *F_ST_* values when compared to *An. cruzii* B (0.46) and *An. cruzii* C (0.62). Another case of sympatry was found in Paranapiacaba – SP, now involving *An. cruzii* A and *An. cruzii* C, with an average *F_ST_* value of 0.39.

**Figure 4.**
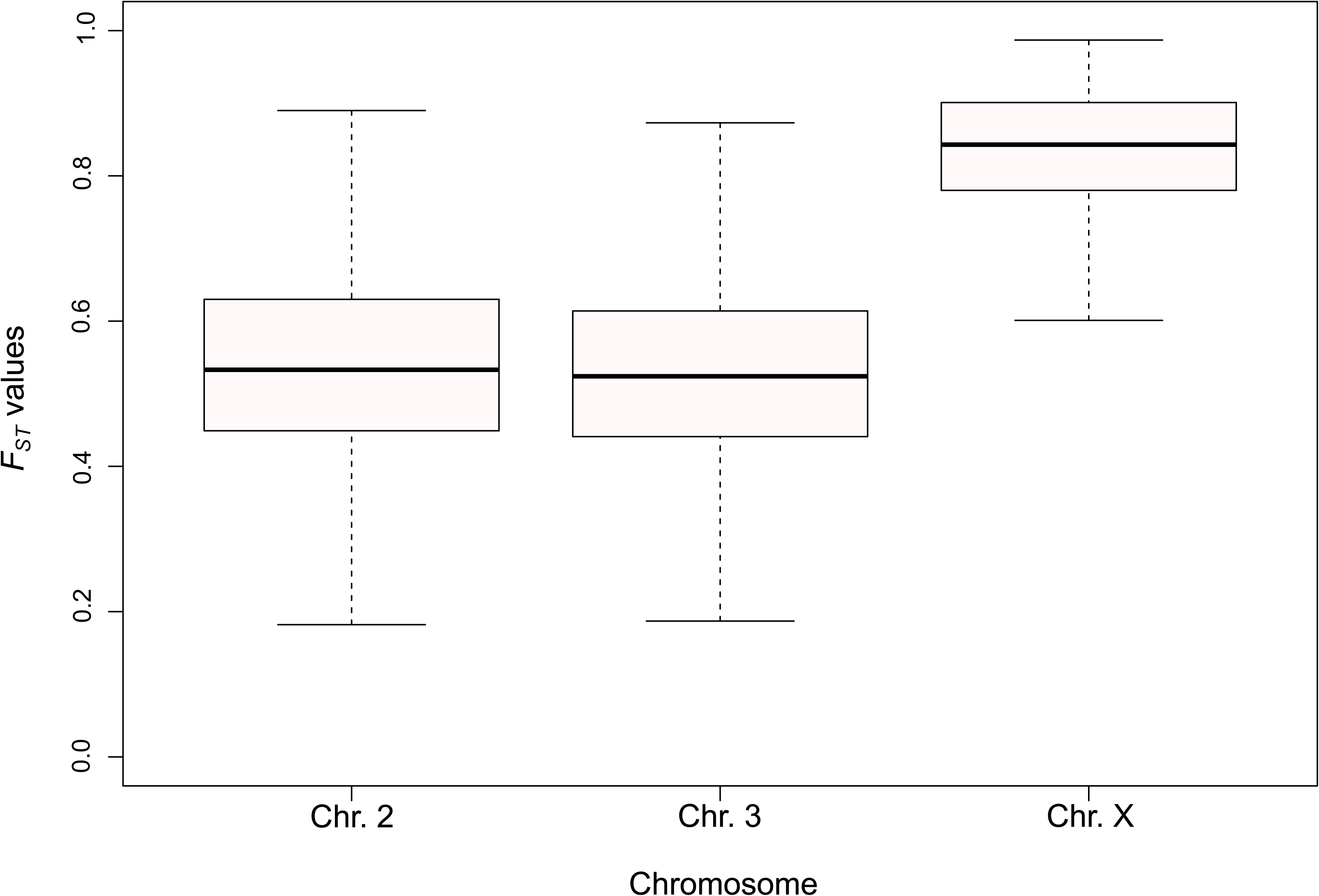
*F_ST_* values by chromosome between *An. cruzii* B *x An. cruzii* C from Bocaina. Note that the *X* chromosome exhibits higher genetic differentiation compared to the autosomes. This pattern suggests that the *X* chromosome plays an important role in the genetic differentiation process of *An. cruzii* s.l. species. The number of genes analysed was 461 for Chr. X, 2,644 for Chr. 2, and 3,558 for Chr. 3.

The highest genetic differentiation among *An. cruzii* s.l. populations was observed in comparisons involving Santa Teresa – ES (*An. cruzii* E; all values close to 0.7), which is the only species with a diagnostic morphological character (see below) (Table 1). The second highest values were observed in comparisons with *An. cruzii* B (*e.g.*, from Bocaina), with mean *F_ST_* values around 0.6 across the autosomes, which exceeds the threshold value (*F_ST_* > 0.35) for species diagnosis proposed by Hey and Pinho^83^. In contrast, comparatively low *F_ST_* values were found between *An. cruzii* C populations (*e.g.*, Bocaina × Campos do Jordão: 0.05; Bocaina × Itatiaia: 0.21; Campos do Jordão × Itatiaia: 0.23), which we deemed as conspecific. Similarly, the comparisons among *An. cruzii* A populations also yielded low *F_ST_* values (*e.g.*, Guapimirim × Paranapiacaba: 0.14; Paranapiacaba × Florianópolis: 0.18; Florianópolis × Guapimirim: 0.21; Table 1, Figure 5), supporting the conclusion that they are a single species. These results are consistent with the positions of these groups in the phylogenomic tree (Figure 3), and in the PCA analysis (see next section).

**Figure 5.**
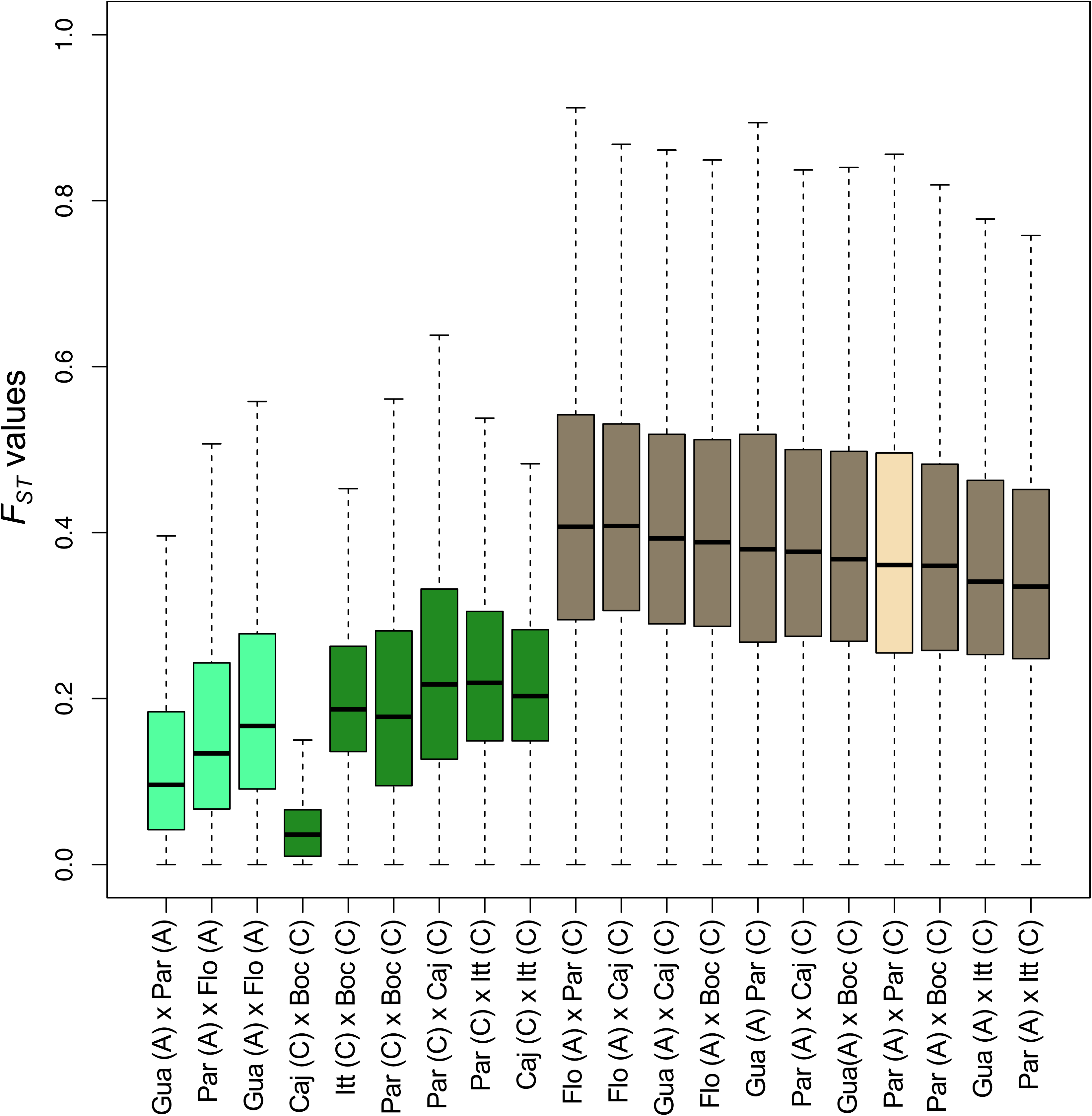
*F_ST_* comparisons within and between *An. cruzii* A and C: intra-, inter-specific, and sympatric *x* allopatric populations. Intra-specific comparisons (green) are contrasted with inter-specific comparisons (brown). Additionally, *F_ST_* values between sympatric (Paranapiacaba A *x* Paranapiacaba C, in cream) and allopatric (brown) populations of *An. cruzii* A and *An. cruzii* C are shown. Note that intra-specific *F_ST_* values (light and dark green) are consistently lower than inter-specific ones (brown and cream). Conversely, the single sympatric inter-specific comparison - Par (A) *x* Par (C) shown in cream - falls approximately in the mid-range of the allopatric comparisons (shown in brown; see Table 1 for exact *F_ST_* values), suggesting that gene flow between these species is very limited or absent. A total of 6,663 genes were analysed. The Y-axis represents *F_ST_* values, and the X-axis indicates the populations in each pairwise comparison. The species designations within the *An. cruzii* s.l. complex (labelled A and C) are shown in brackets next to each population name.

Across all comparisons, the highest *F_ST_* values were detected on the *X* chromosome, suggesting a prominent role for this chromosome in the genetic differentiation process (Figure 4; Table 1; Additional File 5). A similar pattern has been reported for the *An. gambiae* complex^84^, where the *X* chromosome is known to evolve more rapidly, a trend also documented in *Drosophila*, spiders, fish, and other organisms^85–89^.

In addition, we observed high *F_ST_* values for *An. bellator*, for example, between Ilha Grande × Camacan and Camacan × Itaparica, both with mean values around 0.7 (Table 2). These are well above the *F_ST_* > 0.35 threshold for species delimitation proposed by Hey and Pinho^83^, strongly suggesting that *An. bellator* also comprises a cryptic species complex. In contrast, the opposite was observed for *An. homunculus*, with a maximum *F_ST_* of 0.27, even between populations separated by over 1,700 km (Table 3).

**Table 2.** Mean *F_ST_* values of comparisons among *An. bellator* s.l. populations. Number of genes is 6,625 (all), 369 (ChrX), 2,386(Chr1), 3,870 (Chr2). The numbers of individuals for each population are indicated in the parenthesis. Population abbreviations: Ilha Grande (Ilh); Itaparica (Itp); Camacan (Cam).

| Species | Populations | All genes | ChrX | Chr1 | Chr2 |
| --- | --- | --- | --- | --- | --- |
| *An. bellator* A × *An. bellator* B | Ilh (4), Cam (6) | 0.749 | 0.885 | 0.730 | 0.747 |
| *An. bellator* B × *An. bellator* C | Cam (6), Itp (2) | 0.752 | 0.908 | 0.729 | 0.751 |
| *An. bellator* A × *An. bellator* C | Ilh (4), Itp (2) | 0.512 | 0.695 | 0.636 | 0.532 |

**Table 3.** Mean *F_ST_* values of comparisons among *An. homunculus* populations. Number of genes are 6,595 (all), 366 (ChrX), 2,373 (Chr1), 3,856 (Chr2). The numbers of individuals for each population are indicated in the parenthesis. Population abbreviations: Camacan (Cam); Santos (Snt); Santa Teresa (San). Based on the phylogenomic analysis and the *F_ST_* values, all populations were deemed conspecific.

| Species | Populations | All genes | ChrX | Chr1 | Chr2 |
| --- | --- | --- | --- | --- | --- |
| *An. homunculus* | Cam (2), Snt (3) | 0.187 | 0.285 | 0.167 | 0.187 |
| *An. homunculus* | Cam (2), San (2) | 0.225 | 0.328 | 0.118 | 0.225 |
| *An. homunculus* | Cam (2), Flo (1) | 0.175 | 0.337 | 0.210 | 0.175 |
| *An. homunculus* | San (2), Flo (1) | 0.272 | 0.448 | 0.135 | 0.272 |
| *An. homunculus* | Snt (3), Flo (1) | 0.178 | 0.250 | 0.193 | 0.178 |
| *An. homunculus* | Snt (3), San (2) | 0.249 | 0.352 | 0.132 | 0.249 |

### Principal Component Analysis (PCA) and ADMIXTURE

We further analyzed our data using PCA and ADMIXTURE, focusing on the *An. cruzii* s.l. populations because the sample sizes are larger and include both sympatric and allopatric populations. These analyses confirm and extend the previous results.

PCA based on whole-genome SNPs corroborated the phylogenomic analyses, clearly separating *An. cruzii* s.l. specimens into the same five groups (Figure 6a). Regarding ADMIXTURE (Figure 6b; Additional File 6), nearly all *An. cruzii* lineages appeared highly differentiated, with no sign of introgression among them. The exception is *An.cruzii* D, which, for most K values and chromosomes, appeared as an admixture of other lineages, particularly *An. cruzii* A. However, this result may be an artefact: *An.cruzii* D is represented by a single individual, and it is known that ADMIXTURE analyses tend to fit groups containing fewer samples as mixtures of other groups^90^.

**Figure 6.**
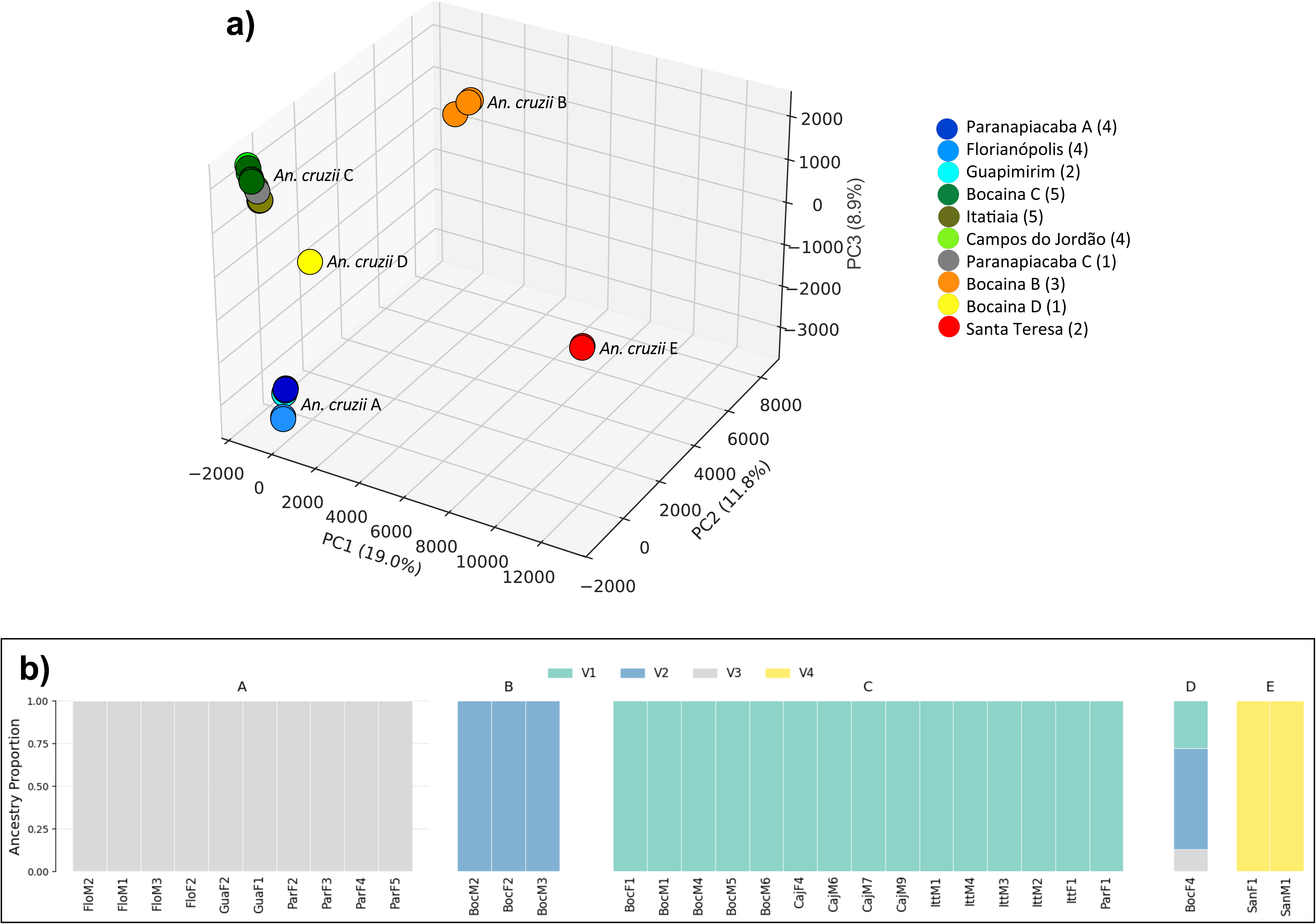
Genomic structure of *An. cruzii* s.l. **a)** Principal component analysis (PCA) based on 20,319,109 biallelic SNPs from the whole genome corroborated the phylogenomic analyses (Figure 3), separating specimens into five groups. **b)** ADMIXTURE analysis for chromosome 2 (K= 4) strongly suggests a lack of recent admixture among the lineages, except for *An. cruzii* D. This result for *An. cruzii D* is not supported at other K values and is probably an artefact (see text). See Additional File 6 for data on other chromosomes and K values.

Another caveat worth mentioning is that ADMIXTURE and related methods assume admixture in the recent history of populations; if there is no admixture, “the algorithm attempts to fit the data as best it can by finding the combination of admixture proportions and ancestral frequencies that best explain the observed patterns”^90^. This may also help explain the odd results we found for *An. cruzii* D. Notably, the very high *F_ST_* values we found among the five *An. cruzii* lineages are more compatible with little, if any, recent admixture. The lack of any clear sign of introgression is an interesting result that contrasts with findings on the *An. gambiae* complex^91,92^. We will return to this point in the Discussion.

### Morphology of the male genitalia

We examined the male genitalia of 16 sequenced samples. Fourteen of these were morphologically classified as *An. cruzii*: three from Florianópolis – SC (*An. cruzii* A), six from Bocaina – SP (two from *An. cruzii* B, four from *An. cruzii* C), four from Itatiaia – RJ (*An. cruzii* C), and one from Santa Teresa – ES (*An. cruzii* E). The only morphological difference observed was in the ventral claspette of *An. cruzii* San M1 from Santa Teresa. Whereas the ventral claspette of *An. cruzii* typically has a lateral expansion that ranges from rounded to sinuous but does not curve posteriorly^38^, in *An. cruzii* San M1 it was neither rounded nor sinuous. Instead, it exhibited a unique golf club-like shape (Additional File 7), which differed significantly from the typical *An. cruzii* ventral claspette and from those of other *Kerteszia* species, including *An. laneanus*^38^. The male genitalia of all other specimens appeared identical, with no discernible morphological differences (Additional File 7). Interestingly, the *An. cruzii* samples from Santa Teresa (*An. cruzii* E) form the most divergent cluster among the *An. cruzii* species in the phylogenetic (Figure 3) and its pairwise *F_ST_* with all other groups are nearly always the highest ones.

The two remaining specimens belonged to *An. bellator*: one from Ilha Grande – RJ (*An. bellator* A) and one from Itaparica – BA (*An. bellator* C). The male genitalia of these two individuals appeared identical. Unfortunately, only female samples from Camacan – BA (*An*. *bellator* B) were available, preventing an analysis of this potentially diagnostic character.

### Chromosomal inversions

One of our Nanopore assemblies, “*An. cruzii* Boc M7 N” (BocM7, for short; Additional File 1), has very high contiguity (largest contig: 34 Mbp; N50: 17 Mbp), allowing us to search for chromosomal inversions by aligning it to the reference genome GCA_943734635.1. These genomes belong to different species: BocM7 was collected in Bocaina – SP and belongs to *An. cruzii* C, whereas the reference genome is from Maquiné – RS and belongs to *An. cruzii* A. Contigs smaller than 1 Mbp were removed from both genomes using *seqtk*^63^, and the remaining contigs were aligned and visualized using *D-GENIES*^93^. Visual inspection revealed two inversions, one in each autosome (Figure 7). The inversion on chromosome 2 spans ∼ 22 Mbp (one-third of the chromosome / approximate coordinates: contig ptg000054l: 5Mbp-27Mbp), while the inversion on chromosome 3 spans ∼ 5 Mbp (approximate coordinates: contig ptg000021l: 3.5Mbp-8.5Mbp).

**Figure 7.**
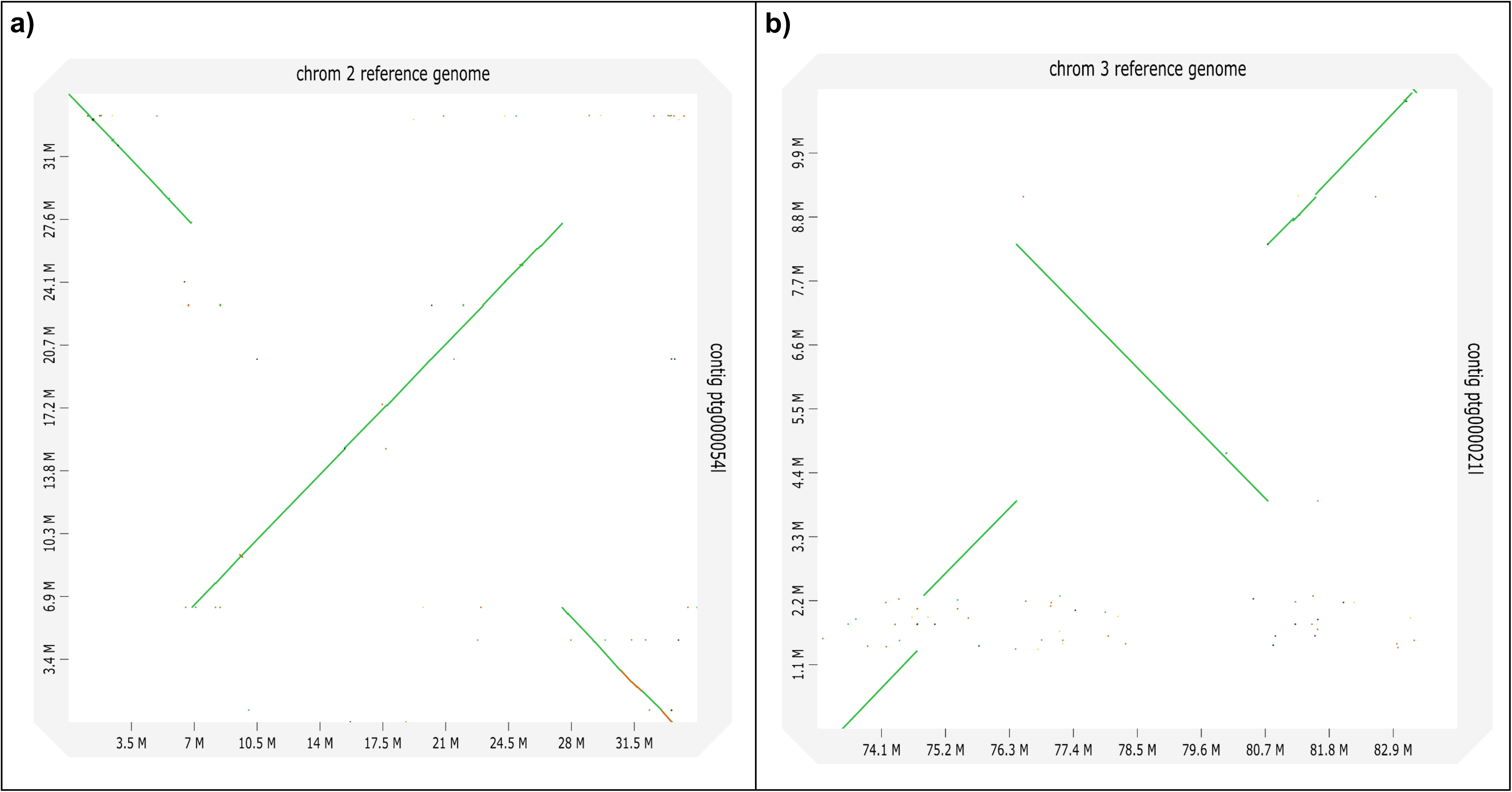
Chromosomal inversions in *An. cruzii* s.l. Dot plots comparing the reference genome of *An. cruzii* A from Maquiné (GCA_943734635.1, X-axis) with the genome of *An. cruzii* C (*An. cruzii* Boc M7 N from Bocaina) (Y-axis), showing two inversions: **(a)** one on chromosome 2 (spanning ∼ 22 Mbp; approximate coordinates: contig ptg000054l: 5Mbp-27Mbp,), and **(b)** another on chromosome 3 (spanning ∼ 5 Mbp; approximate coordinates: contig ptg000021l: 3.5Mbp-8.5Mbp).

It should be noted that additional inversions are likely to be present, since our BocM7 assembly - though highly contiguous - is not at chromosome level. Moreover, the X chromosome is too fragmented in our assembly to allow for visual detection of inversions.

### Divergence times

Divergence times were estimated using a phylogeny constructed with 3,098 orthologous genes from 59 *Kerteszia*. specimens. RelTime^94^ in MEGA 11^74^was applied with four different substitution rates (see Methodology).

This analysis indicates that *An. cruzii* E is the earliest-diverging lineage, dating between approximately 2 and 4 million years ago and showing the greatest divergence time. This is consistent with its more basal position in the phylogeny, its highest *F_ST_* values, and being the only sibling species with a diagnostic difference in male genitalia. *Anopheles cruzii* A (the most widespread species) and *An. cruzii* C (restricted to southeastern mountain regions) diverged more recently, between about 1.2 and 2.5 million years ago (Additional File 8).

## DISCUSSION

The majority of our conclusions regarding the species status of *Kerteszia* populations are based on phylogenomics (*i.e.*, genetic data), so it is worth clarifying the foundations of this type of inference. There are many species concepts and an extensive literature on the subject^95,96^, but we focus here only on the aspects relevant to the present work.

For sexually reproducing organisms, nearly all species concepts agree that if two or more sympatric lineages exhibit strong signals of reproductive isolation (*e.g.*, genome-wide genetic differentiation), then they should be regarded as distinct species. A less clear-cut case arises in the context of allopatry, since some degree of genetic differentiation is expected among conspecific local populations, primarily due to genetic drift and local adaptation^97–99^. According to the biological species concept^100^, in such cases one attempts to infer what would happen if these local populations were to come into contact: if they are expected to merge freely, they are considered conspecific. Conversely, if they are expected to maintain their differentiation due to morphological differences (*e.g.*, in copulatory organs) or high levels of genetic divergence likely to cause mating or developmental incompatibilities (*e.g.*, hybrid sterility), then they are considered separate species.

The fact that many sympatric “good species” exhibit no significant morphological differences highlights the desirability of a genetics-based threshold. But how much genetic differentiation should be adopted as a cut-off to consider allopatric populations as different species? There is no universally accepted answer, but several researchers have approached this question empirically: they have attempted to derive a threshold by analyzing the genetic differentiation of populations that, based on other criteria, had been classified either as conspecific or as belonging to different species^83,101,102^ (but see ref.^103^). In particular, Hey and Pinho^83^, based on a broad range of organisms, proposed that an *F_ST_* value of 0.35 serves as an optimal threshold. However, they also noted considerable overlap between the *F_ST_* distributions of conspecific and non-conspecific populations (see their Figure 6). It should be noted that part of this overlap was undoubtedly due to sampling error, as many of the studies included in Hey and Pinho’s^83^ dataset were based on a small number of loci. We will now interpret our results on *Kerteszia* in light of the framework outlined above.

### Phylogenomic analysis and cryptic speciation in *An. cruzii* s.l

Several studies conducted over the past two decades have provided evidence that *An. cruzii* constitutes a complex of cryptic species^20,22–25,27,104^. These studies were based on one or a few genetic markers, limiting their conclusions’ strength. In the present work, we employed morphology, phylogenomics, and extensive sampling to address this question more robustly.

Our phylogenomic analysis of *An. cruzii* s.l. yielded highly consistent results: the same topology (Figure 3) was recovered using both the multispecies coalescent and concatenation with maximum likelihood approaches, with both stringent (2,480 genes) and relaxed (3,098 genes) datasets. This analysis identified five lineages, provisionally designated *An. cruzii* A – E: *Anopheles cruzii* A is widely distributed along Brazil’s Atlantic coast (Maquiné – RS, Florianópolis – SC, Paranapiacaba – SP, and Guapimirim – RJ); *Anopheles cruzii* B and D were found only in Bocaina – SP; *Anopheles cruzii* C was found across the Serra do Mar and Serra da Mantiqueira ranges (Bocaina – SP, Paranapiacaba – SP, Campos do Jordão – SP, and Itatiaia – RJ); *Anopheles cruzii* E was found exclusively in Santa Teresa – ES (Figure 3).

Figure 8 summarizes the evidence supporting the conclusion that these lineages represent distinct species. *Anopheles cruzii* E displays a morphological difference in the male terminalia that alone supports its recognition as a separate species. Strong genetic differentiation in sympatry at Bocaina provides direct evidence for species status in *An. cruzii* B, *An. cruzii* C, and *An. cruzii* D (*F_ST_* range: 0.47 – 0.62; Table 1), and the same applies to *An. cruzii* A and *An. cruzii* C in Paranapiacaba (*F_ST_*: 0.39).

**Figure 8.**
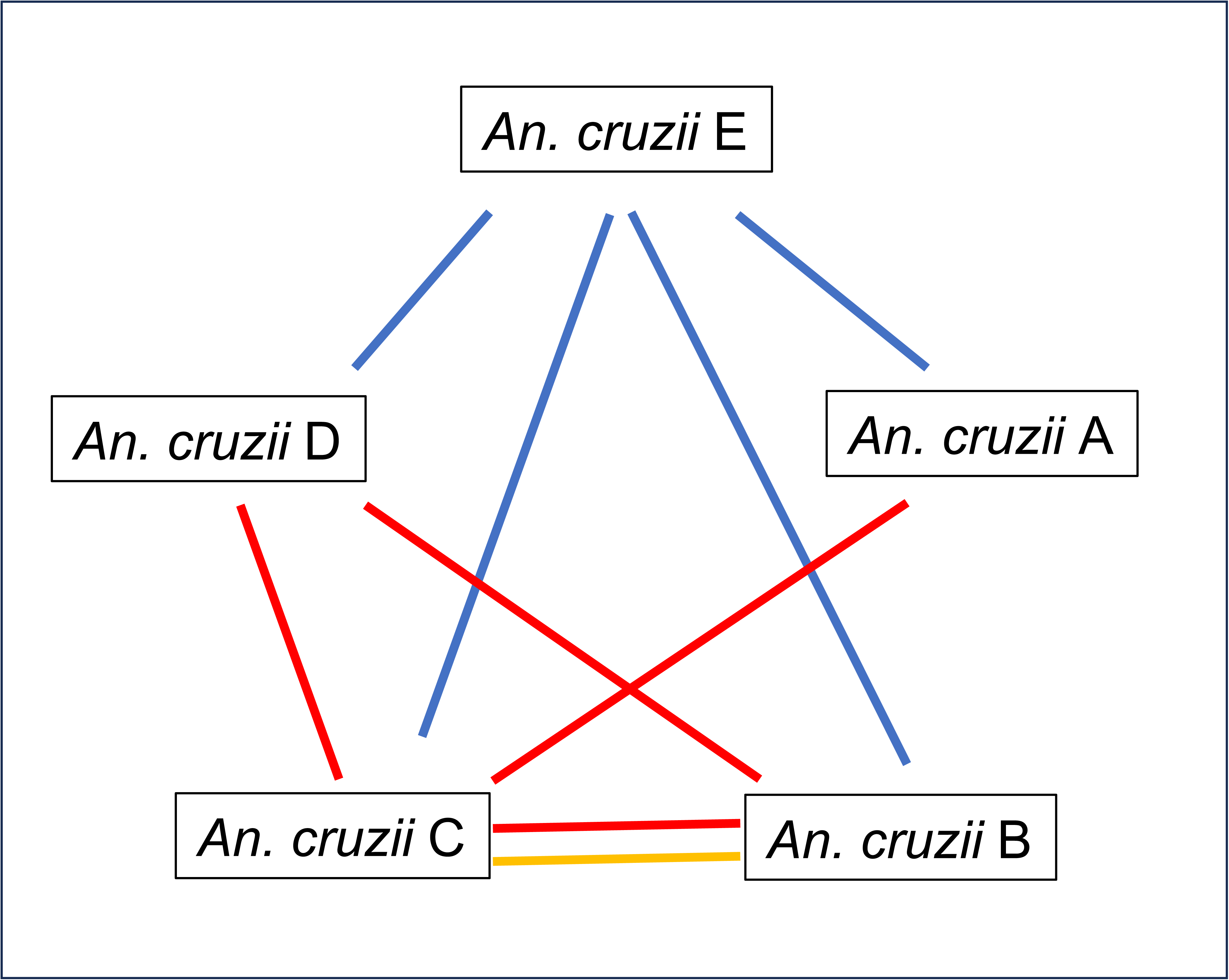
Evidences of cryptic speciation in *An. cruzii* s.l. Taxa connected by red lines have genetically differentiated lineages under sympatry (at the locations of Bocaina and Paranapiacaba), blue lines connect those that have a morphological difference in male genitalia, and the yellow indicates direct evidence of reproductive isolation (lack of heterozygotes in sympatry).

Some species pairs do not co-occur sympatrically in our dataset (*An. cruzii* A × *An. cruzii* B; *An. cruzii* A × *An. cruzii* D), so it is formally possible that they are conspecific. However, their phylogenetic positions (Figure 3) and high *F_ST_* values (range: 0.46 – 0.58; Table 1) strongly support their recognition as distinct species.

Additionally, we genotyped the *cpr* gene in a large sample of wild-caught mosquitoes from Bocaina subsequently identified as *An. cruzii* B and *An. cruzii* C. We detected both homozygotes (*cpr^B^*/*cpr^B^* and *cpr^C^*/*cpr^C^*), but no heterozygotes (*cpr^B^*/*cpr^C^*), strongly suggesting complete reproductive isolation^105^. These patterns satisfy the biological species concept ^95,106^, implying that these lineages represent separate species.

In summary, we believe that the evidence for cryptic speciation within *An. cruzii* is now unequivocal. It is supported by a large number of genetic markers, and we observed two instances of genetically divergent lineages occurring in sympatry (*An. cruzii* B, C, and D in Bocaina – SP, and *An. cruzii* A and C in Paranapiacaba – SP), alongside a diagnostic morphological character (in *An. cruzii* E from Santa Teresa – ES).

Our results suggest that species within *An. cruzii* s.l. diverged between ∼1 and 4 million years ago, consistent with estimates by Rona et al. ^25,26^, who proposed divergence between ∼0.6 and 3.6 Mya. Rona et al.^26^ further hypothesized that Pliocene - Pleistocene climatic fluctuations and Atlantic Forest fragmentation drove this divergence, a mechanism also suggested for amphibians^107,108^ and other mosquitoes^108^. Our divergence times are compatible with this hypothesis.

### Phylogenomic analysis and cryptic speciation in *An. bellator* s.l

The evidence for cryptic speciation in *An. bellator* s.l. is less compelling than in the case of *An. cruzii* s.l., as we did not identify any genetically differentiated lineages occurring in sympatry nor any morphological differences. Nevertheless, we observed deep phylogenetic separation among the three sampled populations and very high *F_ST_* values (range: 0.51 – 0.75; Table 2).

Perhaps the most informative way to interpret these findings is to compare them with the well-supported case of *An. cruzii* s.l. As previously mentioned, the *F_ST_* distributions examined by Hey and Pinho^83^ were based on a wide range of organisms (from plants to mammals) and showed considerable overlap between conspecific local populations and different species – partly due to sampling error, as many studies included very few loci. These factors reduce the confidence in the *F_ST_* = 0.35 threshold they proposed. In contrast, the caveats mentioned above do not apply to our *An. cruzii* s.l. data: (i) the data came from closely related taxa within the *Kerteszia* subgenus; (ii) genetic distances are based on a large number of genes (∼ 6,500; estimated from the VCF files); and (iii) most importantly, as shown in Figure 9, the *F_ST_* distribution is clearly discontinuous, comprising two distinct blocks: one with *F_ST_* ≥ 0.25 (range: 0.05 – 0.25) and another with *F_ST_* ≥ 0.35 (range: 0.37 – 0.73). Independent lines of evidence (genetic differentiation in sympatry; morphology) support the interpretation that the second block corresponds to interspecific comparisons.

**Figure 9.**
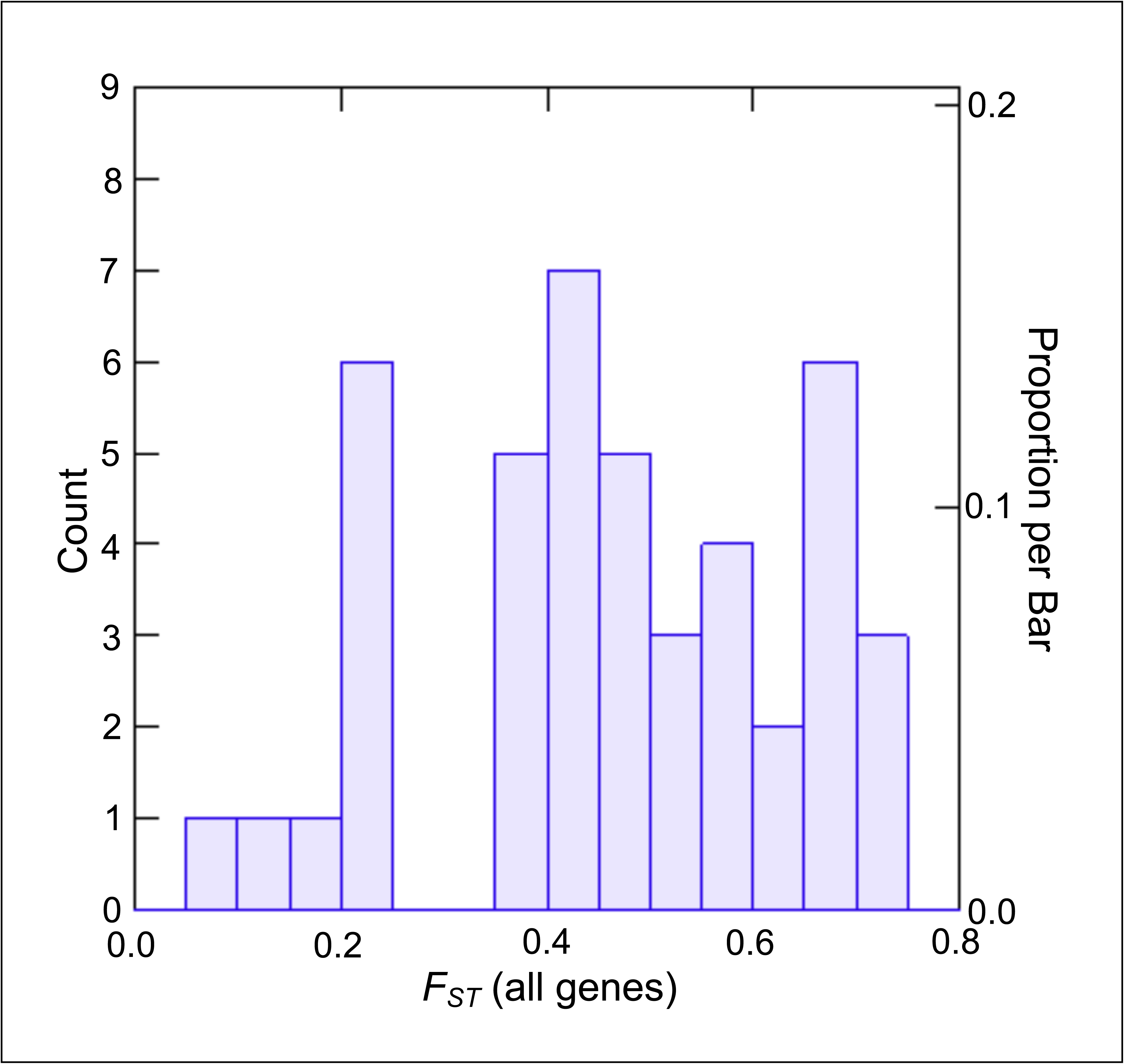
Genetic differentiation among all pairwise comparisons of *An. cruzii* s.l. populations. Note the discontinuity of the distribution: there are two blocks, the first with *F_ST_*<= 0.25 (range: 0.05 – 0.25 and the second with *F_ST_*>= 0.35 (range: 0.37 – 0.73). The left block corresponds to intra-specific comparisons and the right one to inter-specific comparisons (see text).

When placed into the framework shown in Figure 9, the genetic differentiation observed among the three sampled *An. bellator* s.l. populations (*F_ST_* range: 0.51 – 0.75) clearly falls within the interspecific region. Therefore, it is reasonable to conclude that *An. bellator* s.l. constitutes a species complex comprising at least three cryptic species. We provisionally designate these as *An. bellator* A (found in southeastern Brazil: Cananéia – SP and Ilha Grande – RJ), *An. bellator* B (Camacan – BA), and *An. bellator* C (Itaparica – BA). As with *An. cruzii* s.l., geographical distance alone does not explain the observed levels of genetic differentiation: the population from Itaparica – BA is genetically more similar to that from Ilha Grande – RJ (1,300 km apart) than to Camacan – BA (only 300 km apart).

These findings confirm and extend previous studies. Using isoenzymes, Carvalho-Pinto and Lourenço-de-Oliveira^32^ found that *An. bellator* populations from southern and southeastern Brazil (Florianópolis – SC and Cananéia – SP) were genetically similar but distinct from the northeastern population in Bahia State (Itaparica). Using two molecular markers, Voges et al.^33^ identified two *An. bellator* groups in the Brazilian Atlantic Forest: *An. bellator* A, widespread in the southern and southeastern regions (Ilha do Mel – PR, Cananéia – SP, and Ilha Grande – RJ), and *An. bellator* B, found in Camacan – BA.

### Phylogenomic analysis in *An. homunculus*

The case of this species is the opposite of what we observed for *An. bellator* s.l.: the branches in the phylogenetic tree are all very short (Figure 3), and all pairwise *F_ST_* values are low (range: 0.18 – 0.27; Table 3), falling within the intra-specific range observed for *An*. *cruzii* (Figure 9). It is noteworthy that some of these populations are separated by up to 1,700 km. Thus, the available evidence suggests that *An. homunculus* constitutes a single species throughout the Atlantic Forest, consistent with the findings of Cardoso et al.^34^, which were based on very few molecular markers. This conclusion may change if future collections reveal genetically differentiated sympatric lineages, or highly differentiated allopatric populations.

### Are there more species in the *An. cruzii* and *An. bellator* complexes?

The answer is probably yes. First, the logistical challenges of field collection and the cost of genome sequencing inevitably limit sample sizes; additional sampling is likely to uncover additional species (*e.g.*, one of the species we found – *An. cruzii* D – was represented by a single individual). Second, there is the case of *An. laneanus*, which can be distinguished from *An. cruzii* s.l. by subtle morphological features on the tarsus, and by clear differences in the male genitalia^8,109^. Molecular studies based on a small number of markers suggest that it is either part of the *An. cruzii* complex^110^ or forms a sister clade^31,33^. None of the specimens we collected matched the male genitalia of *An. laneanus* (Additional File 7), despite two collection trips to the type locality of this species (Campos do Jordão – SP^111^).

### Comparison of the present results with Dias et al^27^

Dias et al.^27^ studied the same population from Bocaina – SP reported here and were the first to report heterozygote deficiency at the *cpr* gene in *An. cruzii* s.l. They genotyped 12 wild-caught mosquitoes using Sanger sequencing of PCR products and found six *cpr^B^*/*cpr^B^*, five *cpr^C^*/*cpr^C^*, and one heterozygote *cpr^B^*/*cpr^C^*. As reported above, we found three *cpr^B^*/*cpr^B^*, 84 *cpr^C^*/*cpr^C^*, and no heterozygotes. Hence, there are two relevant differences between these collections. First, the frequency of *An. cruzii* B dropped from 54% to 3%, a difference that is statistically significant (*P* < 10^-4^, Fisher’s Exact Test). The collections were conducted in February 2013 (Dias et al.^27^) and February 2023 (present study), so this difference is possibly due to long-term changes in the abundance of the two species.

Regarding the second difference, Dias et al.^27^ found one heterozygote (presumably a hybrid between *An. cruzii* B and *An. cruzii* C), whereas we found none. A proper statistical test for this difference that preserves the statistical power is somewhat tricky due to the significant change in allelic frequency (we would expect a higher frequency of heterozygotes in the 2018 dataset compared to the dataset reported here). We made an approximation by ignoring the difference in allelic frequency, lumping both homozygotes (to increase the statistical power), and then comparing the homozygote *vs* heterozygote counts in the two collections (*i.e.*, we compared 11 homozygotes, 1 heterozygote to 87 homozygotes, 0 heterozygotes). The difference between the collections was not statistically significant (*P* = 0.12, Fisher’s Exact Test). Note that the approximation we used (ignoring the difference in allelic frequencies) is expected to decrease the *P*-value in this case, since we would in any case expect more heterozygotes in the Dias et al.^27^ sample. In other words, the unbiased *P*-value is higher than 0.12. Therefore, we have no evidence that the hybrid frequency changed between the two collections. The main point, however, is that the results of Dias et al.^27^ show that hybrids between *An. cruzii* B and *An. cruzii* C do occur in nature (albeit at low frequency). It should be noted that hybrid formation does not necessary imply gene flow (*e.g*., the hybrids may be completely sterile).

### Comparison between the *An. cruzii* and *An. gambiae* complexes

The pattern we found in *An. cruzii* s.l. resembles, in many aspects, the well-studied case of *An. gambiae* s.l.: both complexes contain cryptic species lacking any diagnostic morphological characters, and in both cases molecular data were essential to identify the biological species^91,92,112–115^. There are, however, two relevant and related differences: *An. cruzii* s.l. species are much more differentiated and show no evidence of gene flow among them. Specifically, in *An. cruzii* s.l., *F_ST_* values are very high and do not appear to decrease in sympatry (Figure 5, Additional File 9), and ADMIXTURE analysis did not reveal any clear sign of introgression in either sympatric or allopatric populations (Figure 6a). These patterns contrast sharply with those observed in species of the *An. gambiae* complex. For example, the smallest pairwise *F_ST_* between different species we found in the *An. cruzii* complex is 0.37 (Table 1), whereas the largest pairwise *F_ST_* reported by Miles *et al*.^92^ for African populations of *An. gambiae* and *An. coluzzii* is 0.14. West African sympatric populations show even smaller *F_ST_* values, ranging from 0 to 0.04^91^. Furthermore, ADMIXTURE analyses clearly indicate introgression within the *An. gambiae* complex^91,92^. It thus appear that the speciation process in the *An. cruzii* lineages we studied is at a much more advanced stage than in the *An. gambiae* complex, and that it is essentially complete.

## CONCLUSIONS AND PERSPECTIVES

We found that *An. cruzii* s.l. comprises at least five cryptic species. One widely distributed group, provisionally labelled *An. cruzii* A, includes populations from the South and Southeast regions of Brazil, in the Serra do Mar coastal zone (Maquiné – RS, Florianópolis – SC, Paranapiacaba – SP, and Guapimirim – RJ). *Anopheles cruzii* B and D were found exclusively in Bocaina – SP. *Anopheles cruzii* C was found in Serra do Mar and Serra da Mantiqueira, two parallel mountain ranges in the Southeast/South of Brazil. (Bocaina – SP, Paranapiacaba – SP, Campos do Jordão – SP, and Itatiaia – RJ). Finally, *An. cruzii* E was found only in Santa Teresa – ES; this is the only species with a diagnostic morphological character. These conclusions are supported, in most cases, by strong genetic divergence in sympatry, and by the aforementioned morphological difference.

Similarly, *An. bellator* s.l. appears to comprise at least three species (provisionally labelled *An. bellator* A, B and C), although in this case the only evidence is the high genetic differentiation observed among allopatric populations.

Finally, *An. homunculus* appears to be a single species, as even populations separated by large geographical distances exhibit only moderate genetic differentiation, within the intra-specific range.

These findings open many promising avenues for research. Genome sequences are now available for all the species mentioned above, in most cases from several individuals. With this resource, it is straightforward to design species-specific PCR primers (as we did for *An. cruzii* B and *An. cruzii* C), enabling a relatively simple and inexpensive method for species identification. This, in turn, will make it possible, for instance, to assess the vector competence for malaria transmission in sympatric species, and to better understand the substantial shift in species composition observed between Dias et al.^27^ and our collection.

## Supporting information

AdditionalFile1

AdditionalFile2

AdditionalFile3

AdditionalFile4

AdditionalFile5

AdditionalFile6

AdditionalFile7

AdditionalFile8

AdditionalFile9

## Competing interests

The authors declare that they have no competing interests.

This paper forms part of the Ph.D. thesis of Kamila Voges, undertaken within the Postgraduate Programme in Cell and Developmental Biology (PPGBCD) at the Center for Biological Sciences (CCB), Federal University of Santa Catarina (UFSC), Brazil.

## Authors’ contributions

KV, LDPR, ABC, CJCP, ANP, GRD and HRR collected the mosquitoes. KV and CJCP conducted the morphological identification. KV, LDPR, ABC, JC, GRD, EGD, FU, SJF, and TV were responsible for data generation, genome assembly, and analysis. KV drafted the manuscript, with critical revisions provided by LDPR, ABC, and GRD. LDPR and ABC contributed to the design and coordination of the study. All authors read and approved the final manuscript.

## Acknowledgements

The authors thank Natália Valério de Souza, Iara Carolini Pinheiro, André Akira Gonzaga Yoshikawa, Sabrina Fernandes Cardoso, João Victor Costa Guesser, Anna Luiza Buainain, Julia Levinstein, Felipe Rocha, and Paulo Paiva for their assistance during fieldwork; Paulo Paiva for critically reading the manuscript; LAMEB – Federal University of Santa Catarina for access to microscopy facilities and technical support; and Dr. Bernard Kim for assistance with Nanopore sequencing.

## ADDITIONAL FILE LEGENDS

**Additional File 1. Detailed information on samples used in this study, including genomic statistics, assembly metrics, and completeness assessments for *Kerteszia* specimens.** *Source*: Bromeliad – immatures collected in the water of bromeliads and reared in the laboratory; CDC + CO_2_ – adults collected using CDC traps with CO_2_; Family – F1 generation of female mosquitoes collected in the field. <u>Genomescope results with k-mer size 21</u>: *Length (GenomeScope)* is the total genome length based on K-mer size; *uniq (%)* is the percent of the genome that is unique (not repetitive); *Hetero. (%)* is the rate of heterozygosity; *Error (%)* is the error rate of the reads; *dup (%)* is the average rate of read duplications; *Coverage* is the genome read coverage. <u>BUSCO v4.1.4 assessed the assemblies using 3,285 conserved</u> <u>Diptera orthologues from the OrthoDB database (odb10;</u> https://busco.ezlab.org/): *Complete (C)* and *Complete single-copy (S)* is the number of genes in each genome with complete sequences; *Complete (D)* is the duplicated genes; *Fragmented (F)* are genes with fragmented sequences; *Missing (M)* are genes absent in each genome. The values in parentheses are the percentage of BUSCOs in each assembly. <u>Assemblies</u> <u>statistics produced with QUAST</u>: *Length* (*assembly*) is the total number of bases in the assembly; *# contigs* is the total number of contigs in the assembly; *Largest contig* is the length of the longest contig in the assembly; *GC (%)* is the total number of G and C nucleotides in the assembly divided by the total length; *N50* is the genome assembly completeness (sequence length of the shortest contig at 50% of the total genome length). Sample ID codes: Flo (Florianópolis), Boc (Bocaina), Cam (Camacan), Caj (Campos do Jordão), Itt (Itatiaia), Gua (Guapimirim), Ilh (Ilha Grande), Par (Paranapiacaba), San (Santa Teresa), Snt (Santos), Itp (Itaparica). Sex and technology codes: M (male), F (female), N (Nanopore). Additional genomes: two genome samples of *An. cruzii s.l.* from Maquiné (GCA_943734635.1, GCA_964417045.1), one *An. cruzii s.l.* from Itatiaia (GCA_964417115.1), and one *An. bellator* from Cananéia (GCA_943735745.1) were obtained from the Sanger *Anopheles* Reference Genomes Project.

**Additional File 2. Busco V3 results comparing SPAdes and Platanus assemblers in seven samples.** Automatic annotation of 2,799 single-copy orthologs from the Diptera reference set of orthologs (odb9). The number of genes with complete sequences (C and S), genes duplicated (D), genes with fragmented sequences (F), or genes absent in each genome (M) is shown in this table. The values in parentheses are the percentage of BUSCOs in each assembly. Flo: Florianópolis; Boc: Bocaina; Itt: Itatiaia; Ilh: Ilha Grande; Itp: Itaparica. M: males; F: females.

**Additional File 3. Results from BUSCO v4.1.4 for samples sequenced with Nanopore.** Automatic annotation of 3,285 conserved orthologous gene assemblies among Diptera from the OrthoDB database (odb10, downloaded from https://BUSCO.ezlab.org/). The number of genes in each genome with complete single-copy and duplicate sequences is shown for each genome in this table. Values in parentheses are the percentage of BUSCOs in each assembly. Statistics of assemblies produced with QUAST. Total Size: Total number of bases in the assembly. # contigs: total number of contigs in the assembly. Longest contig: Length of the longest contig in the assembly. GC (%): total number of G and C nucleotides in the assembly divided by the total length. N50: genome assembly integrity (sequence length of the shortest contig within 50% of the total genome length). Boc: Bocaina; Flo: Florianópolis; F: Female; M: Male; N: Nanopore.

**Additional File 4. Alignment of the *cpr* gene fragment in *An. cruzii* s.l. from Bocaina.** Alignment of the *cpr* gene fragment in *An. cruzii* B (accession numbers: KT724974.1 and KT724976.1 (Dias et al.^27^)), *An. cruzii* C (accession numbers: KT724986.1 and KT724996.1, Dias et al.^27^,, and *An. cruzii* D from Bocaina. The *cpr* sequence from *An. cruzii* D was retrieved from the assembled genome and exhibited several differences compared with the sympatric *An. cruzii* B and C.

**Additional File 5. *F_ST_*-values calculated from orthologous genes used in this analysis.** The *F_ST_* values were calculated for all pairwise comparisons among *An. cruzii* s.l. populations, as well as for *An. bellator* s.l. and *An. homunculus*. Genes from *An. cruzii* were assigned to chromosomes based on the positions of their orthologs in the *An. cruzii* reference genome (GenBank: GCA_943734635.1). For *An. bellator* and *An. homunculus*, chromosomal classification followed the positions of orthologous genes in the *An. bellator* genome (GenBank: GCA_943735745.2), according to the *Anopheles* Reference Genomes Project. Gene ID: Identifier for each gene in the *Anopheles cruzii* genome (GenBank assembly: GCA_943734635.1). Start / End: Genomic coordinates marking the start and end positions of each gene. Population abbreviations: Flo: Florianópolis; Boc: Bocaina; Cam: Camacan; Caj: Campos do Jordão; Itt: Itatiaia; Gua: Guapimirim; Ilh: Ilha Grande; Par: Paranapiacaba; San: Santa Teresa; Snt: Santos; Itp: Itaparica. Taxon abbreviations: Boc A: *An. cruzii* A from Bocaina; Boc B: *An. cruzii* B from Bocaina; Boc C: *An. cruzii* C from Bocaina; ParA: *An. cruzii* A from Paranapiacaba; Par C: *An. cruzii* C from Paranapiacaba.

**Additional File 6. ADMIXTURE results for all chromosomes, and K=4, K=5, and K=6.** No consistent pattern of admixture was found. *Anopheles cruzii* D may appear as an unmixed sample (chromosome X, K=5), or as a composite of the other four species (chromosome X, K=6), which is probably an artefact caused by its small sample size (see text). Overall, the results are consistent with a lack of recent admixture. We ran the same analysis for K values ranging from 2 to 9, which produced qualitatively similar results.

**Additional File 7. Morphological characters of *An. cruzii* s.l. male genitalia.** Male genitalia of four *An. cruzii* s.l. cryptic species. A morphological difference was observed only in the ventral claspette of *An. cruzii* San M1 from Santa Teresa, as indicated by the yellow arrows in **(c)**. This structure differs from the typical *An. cruzii* morphology, taking on a golf club-like shape. The male genitalia of all other specimens were identical, with no detectable morphological differences. **a)** *An. cruzii* Boc M4 (*An. cruzii* B from Bocaina); **b)** *An. cruzii* Boc M2 (*An. cruzii* C from Bocaina); **c)** *An. cruzii* San M1(*An. cruzii* E from Santa Teresa); **d)** *An. cruzii* Flo M2(*An. cruzii* A from Florianópolis).

**Additional File 8. Divergence times estimated using RelTime in MEGA 11.** Divergence times were estimated from a maximum likelihood tree constructed with 3,098 orthologous genes from 59 specimens. RelTime in MEGA 11 was applied with four substitution rates: (i) 19.0 substitutions/kbp/Myr, based on drosophilid colonisation of the Hawaiian Islands^75^; (ii) 16.7 substitutions/kbp/Myr, estimated for drosophilid 3^rd^ codon positions from fossil calibration (Dias et al., *in press*); and (iii) 13.6 and (iv) 27.2 substitutions/kbp/Myr, derived from the spontaneous mutation rate of *Anopheles stephensi* (1.36 ×10-9 substitutions/site/generation)^76^, assuming 10 and 20 generations per year, respectively. This analysis suggests that *An. cruzii* E diverged earliest, between ∼2 - 4 Mya, consistent with its basal phylogenetic position, highest *F_ST_* values, and unique diagnostic difference in male genitalia. *Anopheles cruzii* A (the most widespread species from *An. cruzii* complex) and *An. cruzii* C (restricted to mountain regions from southeast Brazil) diverged more recently, ∼1.2 - 2.5 Mya.

**Additional File 9. *F_ST_* analysis between sympatric and allopatric *An. cruzii* species.** Pairwise *F_ST_* comparisons between sympatric (light colours, bold abbreviations) and allopatric (dark colours) *An. cruzii* species. The sympatric interspecific comparisons - Boc (B) *x* Boc (C), Boc (C) *x* Boc (D), and Par (A) *x* Par (C) - fall in the middle of the range observed for the allopatric comparisons, indicating that gene flow among these species is likely very limited or absent. A total of 6,663 genes were analysed. The Y-axis represents *F_ST_* values, and the X-axis indicates the populations in each pairwise comparison. The species designations within the *An. cruzii* s.l. complex (labelled A, B, C,and D) are shown in brackets next to each population name. Flo: Florianópolis; Boc: Bocaina; Caj: Campos do Jordão; Itt: Itatiaia; Gua: Guapimirim; Par: Paranapiacaba; San: Santa Teresa.

