## AdditionalFile4 for "*Anopheles (Kerteszia) cruzii*, the main malaria vector in the Brazilian Atlantic Forest, is a complex of at least five cryptic species"

```
KT724974.1        GTGTAATATGGTAAGCGAA-CG----AGAGAGAGTCCTCGCCTATACGGTGACG
KT724976.1        GTGTAATATGGTAAGCGAA-CG----AGAGAGAGTCCTCGCCTATACGGTGACG
KT724986.1        GTGTAATATGGTAAGCGAAACGAGAGGGAGAGAGTCCGCGCCTATACGGTGACG
KT724996.1        GTGTAATATGGTAAGCGAAACGAGAGAGAGAGAGTCCGCGCCTATACGGTGACG
An_cruzii_D       GTGTAATATGGTAAGCGAAA--AGAGAGAGAGAGTCCGCGCCTATACGGTGACG
                  *******************        ********** ****************

KT724974.1        CCGGCGGG----CGCGCCAGCATGTTGTAATCCGTTCCGTTCCACTCTCTCTCT
KT724976.1        CCGGCGGG----CGCGCCAGCATGTTGTAATCCGTTCCGTTCCACTCTCTCTCT
KT724986.1        CCGGCGGGCGGGCGGGCCAGCATGTTGTAATCCGTTCCGTTCCAC------TCT
KT724996.1        CCGGCGGGCGGGCGAGCCAGCATGTTGTAATCCGTTCCATTCCAC------TCT
An_cruzii_D       CCGGCGGG----CGGGCCAGCATGTTGTAATCCGTTCCGTTCCAC------TCT
                  ********    ** *********************** ******      ***

KT724974.1        CTCTGTGCACACGCAGGAAGAGCTGTTGCAGCTGAAAGACATCGAGAAATC
KT724976.1        CTCTGTGCACACGCAGGAAGAGCTGTTGCAGCTGAAAGACATCGAGAAATC
KT724986.1        CTCTGTGCACACGCAGGAAGAGCTGTTGCAGCTGAAAGACATCGAGAAATC
KT724996.1        CTCTGTGCACACGCAGGAAGAGCTGTTGCAGCTGAAAGACATCGAGAAATC
An_cruzii_D       CTCTGTGCACACGCAGGAAGAGCTGTTGCAGCTGAAAGACATCGAGAAATC
                  ***************************************************
```
