## Supplementary figures and images for "*Anopheles (Kerteszia) cruzii*, the main malaria vector in the Brazilian Atlantic Forest, is a complex of at least five cryptic species"

### AdditionalFile6

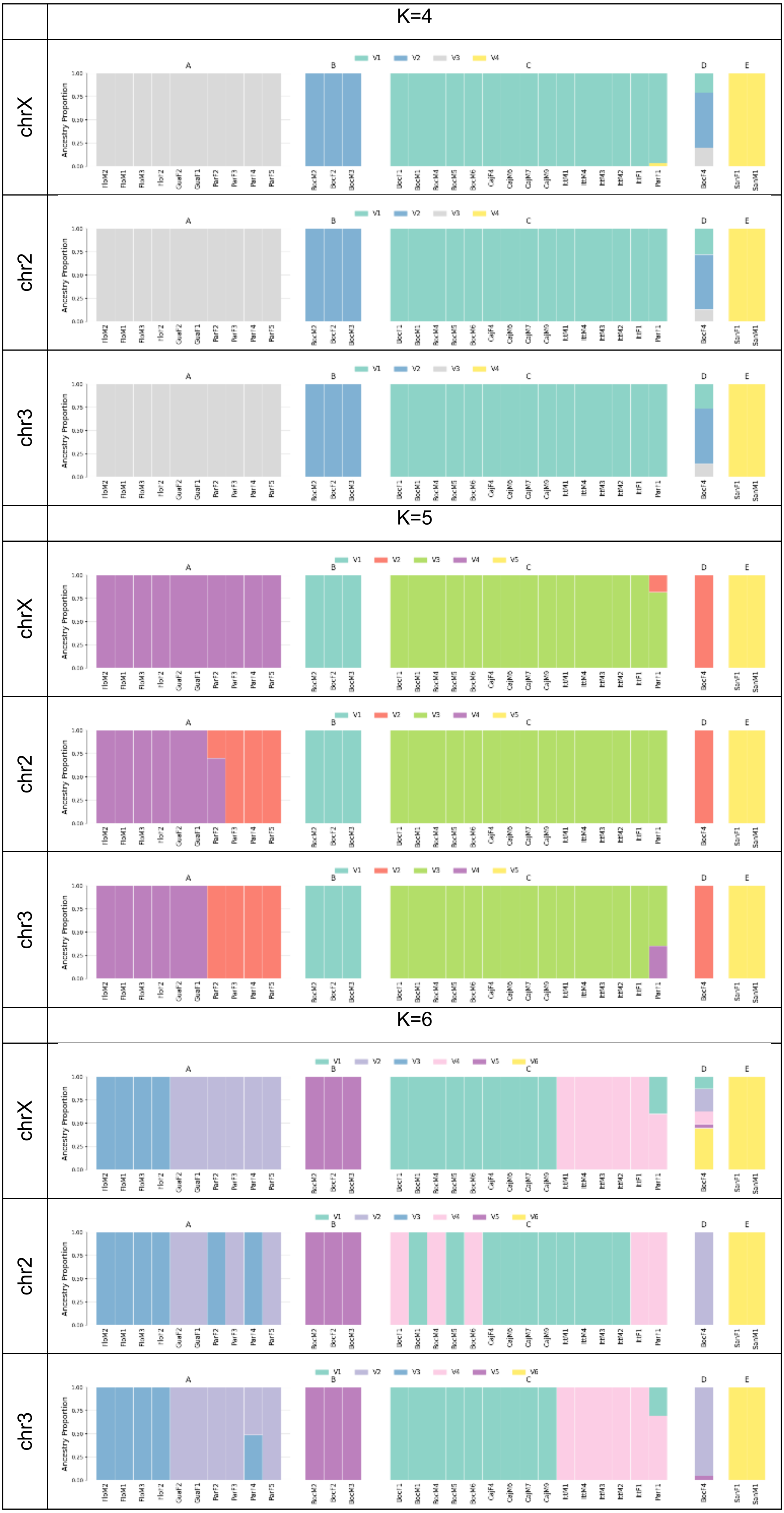

### AdditionalFile7

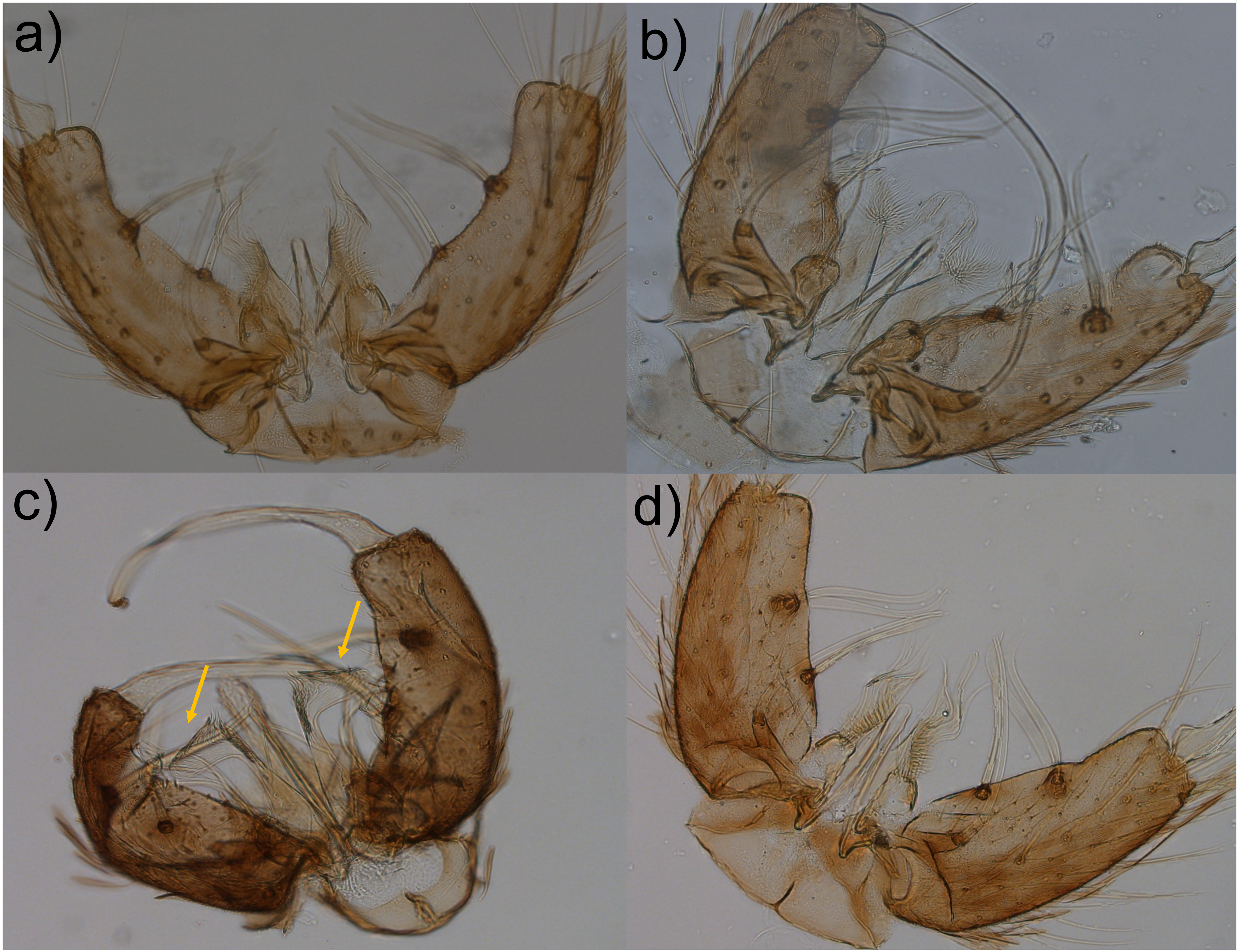

### AdditionalFile9

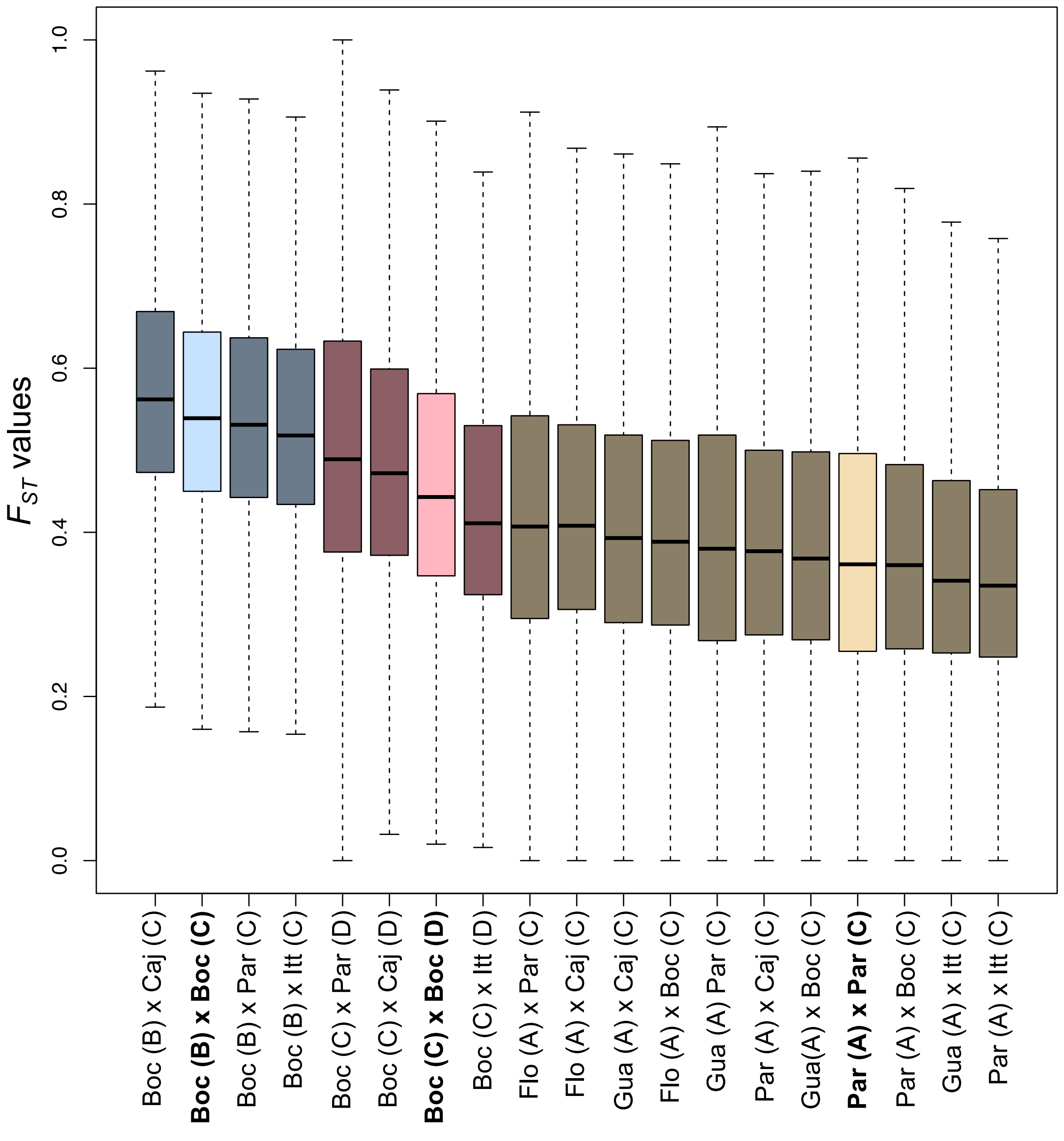
