## AdditionalFile8 for "*Anopheles (Kerteszia) cruzii*, the main malaria vector in the Brazilian Atlantic Forest, is a complex of at least five cryptic species"

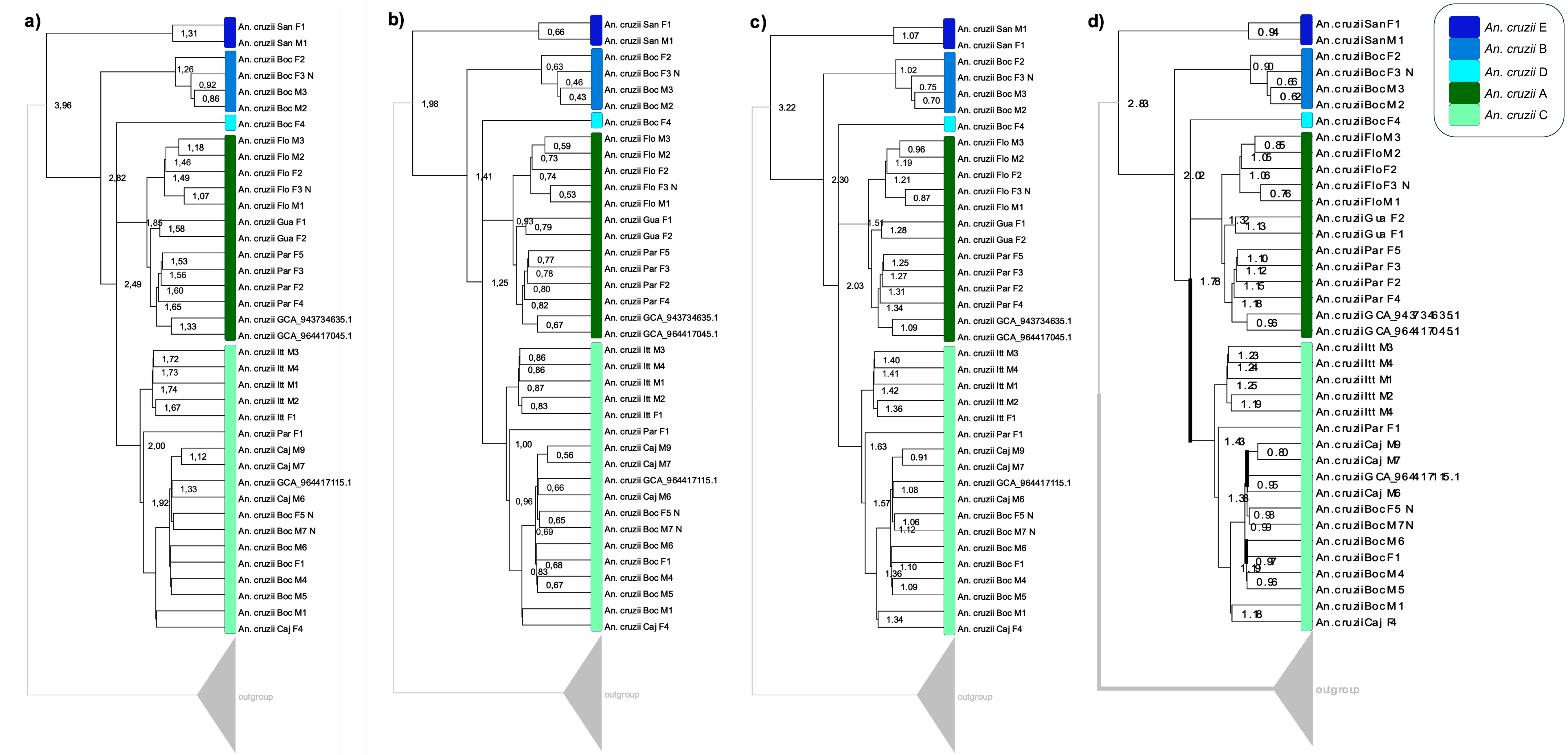


*Anopheles stephensi*
Rashid et al. (2022), 10 generations

*Anopheles stephensi*
Rashid et al. (2022), 20 generations

*Drosophilidae*
Dias et al. (*in press*)

*Drosophila*
Obbard et al. (2012)
